# Discrete dynamic model of the mammalian sperm acrosome reaction: the influence of acrosomal pH and biochemical heterogeneity

**DOI:** 10.1101/2021.03.07.434290

**Authors:** Andrés Aldana, Jorge Carneiro, Gustavo Martínez-Mekler, Alberto Darszon

## Abstract

The acrosome reaction (AR) is an exocytotic process essential for mammalian fertilization. It involves diverse biochemical and physiological changes that culminate in the release of the acrosomal content to the extracellular medium as well as a reorganization of the plasma membrane (PM) that allows sperm to interact and fuse with the egg. In spite of many efforts, there are still important pending questions regarding the molecular mechanism regulating the AR. Particularly, the contribution of acrosomal alkalinization to AR triggering in physiological conditions is not well understood. Also, the dependence of the proportion of sperm capable of undergoing AR on the biochemical heterogeneity within a sperm population has not been studied. Here we present a discrete mathematical model for the human sperm AR, based on the biophysical and biochemical interactions among some of the main components of this complex exocytotic process. We show that this model can qualitatively reproduce diverse experimental results, and that it can be used to analyze how acrosomal pH (pH_*a*_) and cell heterogeneity regulate AR. Our results confirm that pH_*a*_ increase can on its own trigger AR in a subpopulation of sperm, and furthermore, it indicates that this is a necessary step to trigger acrosomal exocytosis through progesterone, a known physiological inducer of AR. Most importantly, we show that the proportion of sperm undergoing AR is directly related to the detailed structure of the population biochemical heterogeneity.

## 1 Introduction

The Acrosome reaction (AR) is an exocytotic process in sperm that is essential for fertilization in many species, including mammals. It involves multiple and complex biochemical and physiological changes that culminate in the release of different hydrolytic enzymes to the extracellular medium, as well as reshaping of the plasma membrane (PM).

The AR confers sperm the ability to interact and fuse with the egg [9, 44, 45], but despite many years of research, it is still to be completely elucidated what are the physiological stimuli of this fundamental process. Furthermore, there are pending questions about for which species the hydrolytic enzymes released to the extracellular medium facilitate the passage of sperm through the zona pellucida (ZP) [23, 32, 45, 72, 101]. The specific biochemical and physiological events that trigger the AR remain elusive, although we know that in the course of this process there is an increase in the intracellular concentration of Ca^2+^ ([Ca^2+^]_*i*_) [23, 80, 88] as well as an increase in intracellular pH (pH_*i*_) [70, 88]. The regulation of pH_*i*_ is important for the functioning of a variety of proteins. The sperm exclusive Ca^2+^ channel (CatSper) and K^+^ channel (Slo3) are strongly pH_*i*_ dependent [16, 70, 105]. The rise in intra acrosomal pH (pH_*a*_) can lead to increases in [Ca^2+^]_*i*_ and spontaneous AR [15, 69] both in mouse and human sperm [15]. However, the relevance of acrosomal alkalinization under a physiologically triggered exocytotic process is not well understood. It is known that Pg promotes [Ca^2+^]_*i*_ elevation by stimulating CatSper in human sperm [4, 64].

Cell population studies indicate that only a fraction of sperm is capable of undergoing AR, either spontaneously or after induction with progesterone (Pg), a known AR inducer present in the female tract at the relevant concentration. In human and mice sperm samples, 15-20% of cells undergo spontaneous AR[69], whereas only 20-30% undergo Pg induced AR[88]. Although this suggests that biochemical heterogeneity plays a role in determining the proportion of sperm capable of undergoing AR either spontaneously or after Pg induction, such heterogeneity has not yet been studied. Moreover, the AR develops progressively in time [80], implying that at a particular time, sperm are heterogeneous in their biochemical states. This hypothesis is supported by reports in the literature showing a wide non uniform range of values for the concentration of distinct intracellular components in a sperm population for different species [2, 55, 65].

In the present work we implement a generalization of the Gene Regulatory Network as formalized by Stuart Kauffman in 1969 [47] and used in many different systems [29, 60, 61], to construct a mathematical and computational model that represents the main biochemical interactions involved in AR. We show that this model can qualitatively reproduce many of the experimental results reported in literature and we use it to characterize how the biochemical heterogeneity in a sperm population affects the proportion of cells capable of displaying spontaneous and Pg-induced AR.

Our model corroborates that acrosomal alkalinization can trigger AR by itself in a fraction of the sperm population and suggests it is important for AR induction by Pg in another fraction. Together, our results indicate that biochemical heterogeneity is closely related with the proportion of sperm capable of displaying AR, and that a pH_*a*_ increase is an essential event in the process of the AR.

## 2 Regulatory network background

### 2.1 AR Preconditions

Capacitation is a precondition necessary for physiological AR in mammalian sperm. Capacitation is a complex process involving plasma membrane remodeling, cholesterol removal, extensive changes in protein phosphorylation patterns and increases in pH_*i*_ and [Ca^2+^]_*i*_ [9], as well as membrane hyperpolarization [12, 27, 88]. Only a subpopulation of sperm (20-40%) becomes capacitated, and the mechanisms involved in such selective capacitation and in eliciting the aforementioned cellular changes are far from clear.

Cell pH_*i*_ regulation is performed mainly by H^+^ fluxes between the extracellular medium, the cytosol and internal stores, as well as HCO_3_^−^ transport and metabolism [2, 13, 40, 53, 69, 70, 78, 84]. In the plasma membrane a Na^+^/HCO_3_^−^ cotransporter (NBC) allows HCO_3_^−^ uptake [70, 78] while a pH dependent Cl^−^/HCO_3_^−^ exchanger (SLC) extrudes it to the extracellular space. Increases in HCO_3_^−^ intracellular concentration ([HCO_3_^−^]_*i*_) also activate a soluble Adenilate Cyclase sAC [17, 49, 70, 73]. The sperm specific Na^+^/H^+^ exchanger (sNHE) contributes importantly to cytosolic pH_*i*_ regulation in mouse sperm [69, 70, 99], though recent results indicate it is NHA_1_ that is important for the ZP induced mouse sperm pH increase during AR [3]. In human sperm the *H*^+^ channel Hv1, appears to be the main pH_*i*_ regulator [13, 28, 53, 63, 70].

The acidic intra acrosomal space (pH_*a*_ ~ 5.4) is maintained principally by a H^+^ V-ATPase in the acrosomal membrane [15, 69, 90]. In turn, the acidic pH_*a*_ contributes to cytosolic acidification by means of a nonspecific acrosomal pH_*a*_ dependent H^+^ outward current (HLeak_*a*_) [15, 69]. It has been proposed that a somatic Na^+^/H^+^ exchanger in the acrosome membrane could participate in this flux [68, 71], however its existence has not been established. During capacitation pH_*a*_ increases and this elevates spontaneous AR [69]. Because of this, it was proposed that an increase in pH_*a*_ during capacitation may be a requirement to prepare sperm for the AR. In this direction, permeable weak bases able to alkalize the acrosome can release Ca^2+^ from acidic compartments including the acrosome, increase [Ca^2+^]_*i*_ and induce AR [15]. On the other hand, it has been proposed that acrosomal alkalinization is involved also in acrosome swelling and outer acrosomal membrane(OAM) deformation during AR by means of pH_*a*_ dependent activation of proteolytic enzymes that destabilize the acrosomal matrix [15, 37].

#### Membrane potential regulation

There is evidence that membrane hyperpolarization is important for capacitation and therefore necessary for AR to occur. Also, a depolarization in capacitated sperm can modulate AR [24, 27]. These events suggest the existence of a fine membrane potential (Em) regulation during capacitation and AR. Em regulates and is regulated by different ionic transporters. Membrane depolarization activates voltage dependent channels like classical Ca_v_s [23, 104], a controverted subject, CatSper [16], K^+^ channels (IKsper) [16, 105] and Hv1 [13, 53]. Em also influences sNHE activity (in mouse sperm) that is activated by hyperpolarization and participates in sperm alkalinization [16]. Cationic efflux carried by IKsper and Hv1 promote hyperpolarization, as well as HCO_3_^−^ entry by NBC activation. Finally, Ca^2+^ uptake through CatSper and possibly by Ca_v_s, and store operated Ca^2+^ channels (SOCs), aside from increasing [Ca^2+^]_*i*_ depolarize Em [18, 24, 86].

### 2.2 The Acrosome Reaction

The acrosome is an acidic vesicle of lysosomal/Golgi origin located at the apical part of the sperm head that accumulates Ca^2+^ in its interior. It contains hydrolytic enzymes released through exocytosis to the extracellular medium when the AR occurs [20, 21, 24]. Although the specific sequence of events that trigger the AR after capacitation is not fully understood, there is evidence that it involves elevations of pH_*i*_ and [Ca^2+^]_*i*_, and acrosomal Ca^2+^ release [24]. These events promote and culminate in acrosome swelling, deformation of the outer acrosomal membrane (OAM), interaction and docking with the plasma membrane (PM) and finally, fusion between OAM and PM that promotes exocytosis [58, 85].

#### Cytosolic and acrosomal Ca^2+^ regulation

It is well established that orchestrated [Ca^2+^]_*i*_ elevations are crucial for the AR. In the sperm head, upon AR induction, Ca^2+^ enters the cytosol mainly through SOCs in the PM, as indicated by indirect evidence [86]. Release of inositol 3-phosphate (IP_3_) caused by phospholipase C delta 4 (PLC_*δ*_) activates IP_3_ receptors (IP_3_R) in the OAM [24, 35, 36, 58, 86, 93]. The latter is a [Ca^2+^]_*i*_ and IP_3_ dependent Ca^2+^ channel with one binding site to IP_3_ and two binding sites to Ca^2+^ of low and high affinity that promote opening or inactivation of the channel, respectively [7, 8, 34, 81]. In the presence of IP_3_, a moderate increase in [Ca^2+^]_*i*_ results from IP_3_ receptors (IP_3_R) opening while high concentrations block the channel. IP_3_R activation releases Ca^2+^ in the acrosome. This event is detected by stromal interaction molecule (STIM) proteins in the OAM [24, 85] which in turn reorganizes and activates SOCs in the PM [51, 86]. In the flagellum, CatSper, a pH dependent and mildly voltage gated Ca^2+^ channel increases [Ca^2+^]_*i*_ concentration [48, 54]. Two different types of Ca^2+^ ATPases help controlling Ca^2+^ elevation in the cytosol promoted by SOCs and IP_3_Rs. The plasma membrane Ca^2+^ ATPase (PMCA) in the PM extrudes Ca^2+^ to the extracellular medium [19, 82] and an acrosomal Ca^2+^ ATPase (ACA) contributes to maintain the high level of [Ca^2+^]_*a*_ during the final stages of the AR. There is debate on which type of Ca^2+^ ATPase is contributing to Ca^2+^ mobilization from the acrosome. The secretory pathway Ca^2+^ ATPase pump type 1 (SPCA1) is expressed in human sperm, although it seems to be localized mainly in the neck region [38]. Also, a different study shows that the sarco/endoplasmic reticulum Ca^2+^ ATPase type 2 (SERCA2) is expressed in the acrosome and mid piece region [50]. The contribution and presence of these two forms of Ca^2+^ ATPase in the AR are yet to be fully elucidated. Ca^2+^ modulates diverse signaling pathways: activation of PLC that produces diacylglycerol (DAG) and IP_3_ [24, 67], activation of AC that increases cyclic adenosine monophosphate (cAMP) levels as well as phosphodiesterases (PDE) that create the opposite effects [46, 86], and activation of synaptotagmin (SYT) [41], a Ca^2+^ sensor important in the final stages of membrane fusion [58].

#### Triggering Membrane Fusion

Fusion between OAM and MP during AR has not been fully characterized. However, several proteins involved in other exocytotic processes have been found in sperm, such as the Ras related protein Rab3A [43, 100, 103], N-ethylmaleimide-sensitive factor (NSF) [62, 76], α - soluble NSF Attachment Protein (αSNAP) [94], SNAP receptor family members (SNARE’s) [75, 83, 95], SNARE’s associated proteins like complexin [77, 106], Ca^2+^ regulated proteins like SYT [41, 42, 75] and calmodulin [96] as well as cellular transport-related proteins like dynamin [58, 106]. Based on multiple literature reports, a model has been developed to describe the specific biochemical events involved in membrane fusion during AR [58]. This model is also consistent with experimental observations in human sperm and we take this as a starting hypothesis for our work. It considers that at early fusion stages SYT is inactive and there are inactive *cis*-SNARE complexes assembled between the OAM and PM which keep the AR in standby until other events trigger it. Further in time IP_3_R channels open and release [Ca^2+^]_*a*_ from the acrosome which promotes SOC channel aperture and an increase in [Ca^2+^]_*i*_. This stimulates the activity of soluble and/or transmembrane AC increasing cAMP levels [5, 11, 87]. There is evidence that cAMP promotes deformation and swelling of the acrosome, possibly activating Rab3A through exchange proteins directly stimulated by the rise in this messenger (EPAC), given the rise in cAMP concentration [58, 102]. The events connecting [Ca^2+^]_*i*_ with cAMP, EPAC and Rab3A are not clear but we consider in our network the proposal that after activation of Rab3A, NFS and αSNAP, *cis*-SNARE complexes disassemble and reorganize in *trans*-SNARE complexes [26]. At this stage SYT dephosphorylates and fusion stands by until Ca^2+^ is released again from the acrosome to activate it. The model hypothesizes that the free Ca^2+^ in the cytosol is inaccessible to the small space between the OAM and the PM at the final stages of the AR, and that the opening of local IP_3_Rs provide the necessary Ca^2+^ that activates SYT and thus SNARE’s, finally allowing the fusion between membranes.

### 2.3 Regulatory Network implementation

The components and interactions of the signaling pathway involved in acrosomal reaction in human sperm, described in the previous section, are illustrated in Figure (1). This signaling pathway was translated formally into the *Regulatory Network* represented in Figure (2), consisting of 38 nodes and 80 links between nodes. In this network, each node represents one of the main components of the AR signaling pathway and links represent regulatory interactions among components. Given any two nodes A and B, their interactions can be positive, negative or dual depending on whether the activity of A promotes (black arrows) or inhibits (red arrows) the activity of B, or its action is context dependent (yellow arrow) on the activity of B. The network includes four *input* nodes that represent external stimuli, namely Ca^2+^ ionophore A23187, progesterone (Pg) and external concentrations of Ca^2+^ and HCO_3_^−^. An *output reporter* node (Fusion or F) representing the fusion between the outer acrosomal membrane and the plasma membrane, indicates the completion of the AR and is the end point of the network.

**Figure 1:**
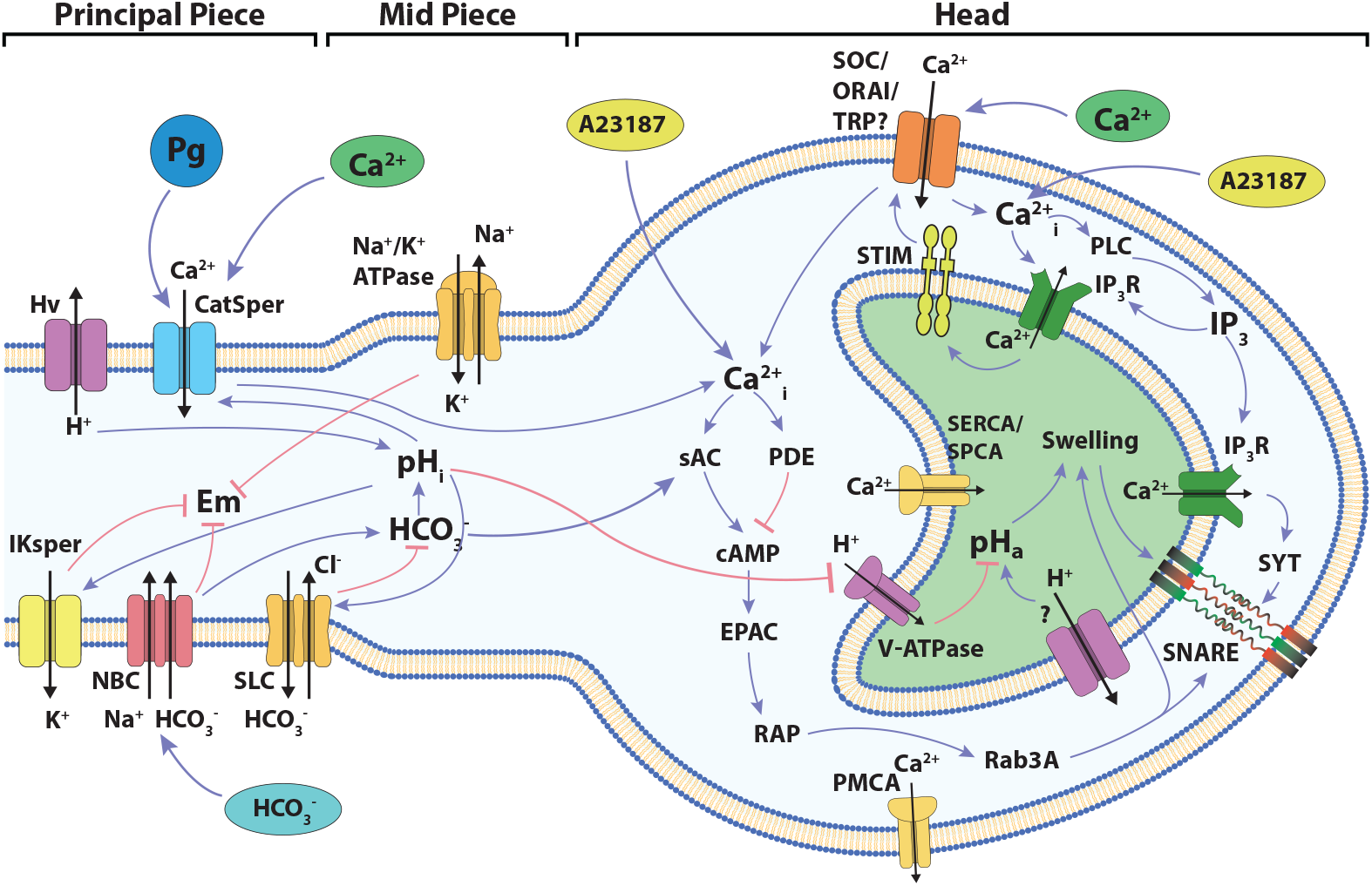
Diagram of the signaling network involved in the development of acrosome reaction in human sperm. The cartoon shows the main components and signaling events to be considered in the development of the model described in the present work. Blue links connecting components indicate positive interactions while red links indicate negative interactions. The localization of the different components among the head and the flagellum are indicated.

**Figure 2:**
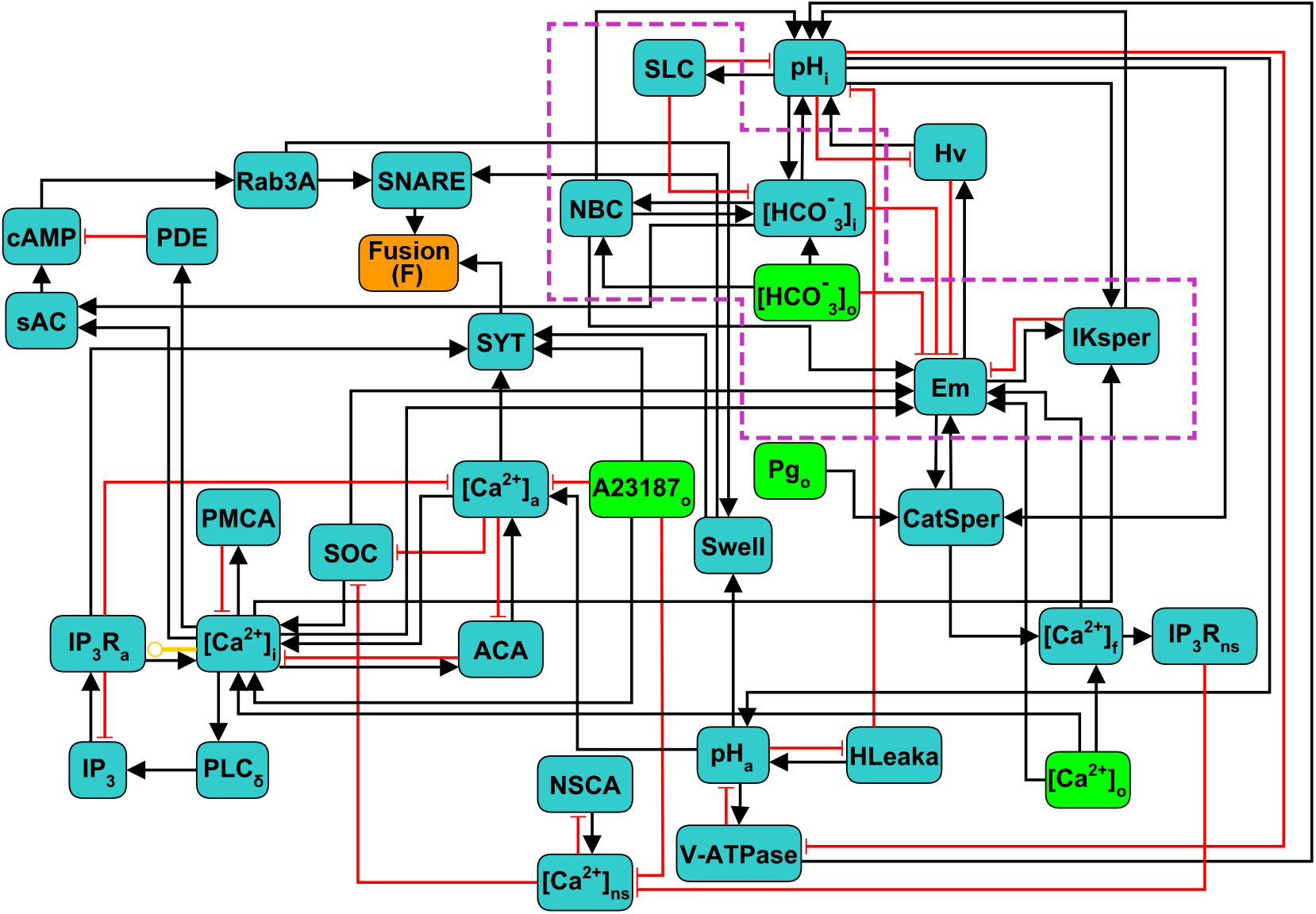
Scheme of the discrete regulatory network for AR in human sperm. Light green nodes indicate extracellular components that can be considered as external stimuli or inputs of the network, dark green nodes represent intracellular components, the orange node Fusion (F) is the network’s output reporter, representing the fusion between OAM and PM, that indicates AR completion. Black arrows indicate positive or activating interactions, red arrows indicate negative or inhibitory interactions, yellow arrow indicates a dual interaction, depending on the values of the nodes involved. Subscripts indicate the physical location in the sperm of the biochemical component represented by its corresponding node: i,a,o,f and ns represent intracellular, acrosomal, outer (extracellular medium), flagellar and neck store respectively. Nodes inside the purple doted box are mostly involved in capacitation and are included as control variables of the necessary conditions to undergo AR.

To describe and analyse the dynamical properties of this model, we use a generalization of the Boolean gene regulatory networks proposed by Stuart Kauffman in 1969 [47] that have been successfully used in different gene regulation studies [60, 61, 47], as well as in biochemical regulation [14, 29, 92]. In this model, the state of the whole network is described by a set of N discrete variables *x_1_,x_2_, …, x_N_*, each one representing the state of one node. Most of the nodes can take values 0 or 1, corresponding to the basal and increased activity of the component, respectively. To better describe the activity and dynamics of some components, 4 nodes take more than 2 discrete values from 0, 1, 2 or 3 as shown in Table (1): Em, CatSper, [Ca^2+^]_*i*_ and pH_*i*_. The particular value of these nodes has different interpretations for each case: Em (hyperpolarized 0, equilibrium 1, mildly depolarized 2, and fully depolarized 3); CatSper (closed 0, open 1, inactivated 2); [Ca^2+^]_*i*_ (basal 0, activator 1, inhibitor 2); pH_*i*_ (acidic 0, mildly alkaline 1, fully alkaline 2). The *node state* is determined by a *regulatory function* that takes as argument the values at a particular time of its regulator nodes. Let us define *x_i1_(t), x_i2_(t),…, x_ik_(t)*, the values at time *t* of the *k* regulators of node *x_i_*. Then the value of node *x_i_* changes at each time step according to:

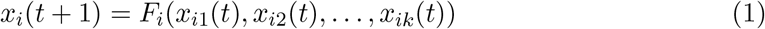

**Table 1:**
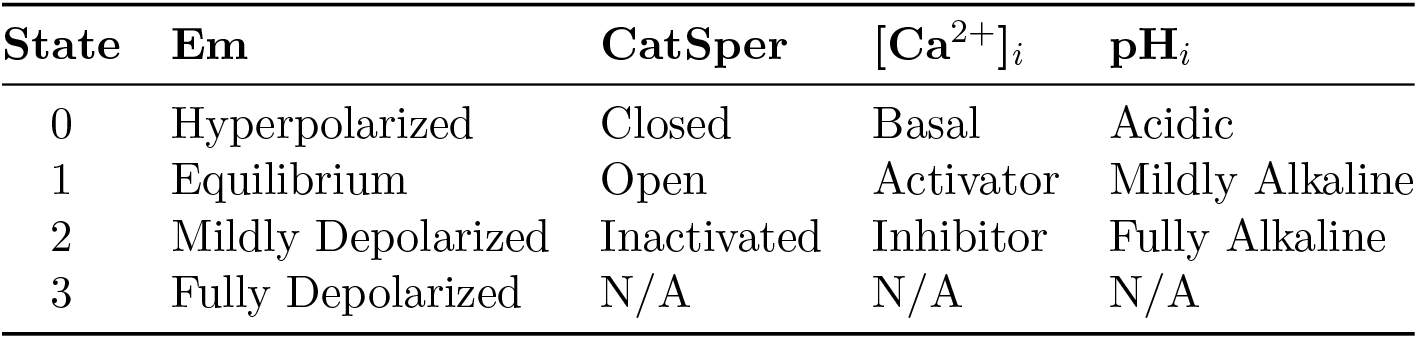
States and interpretation of the non binary nodes. The number of states and their biological interpretation depends on how each node regulates and is regulated by other nodes. The nodes with value N/A (Not Applicable) in state 3 are ternary and only can take states in {0,1,2}.

Where *F_i_* is the regulatory function of node *x_i_*. Since all nodes *x_i_* are updated at each time step, this induces deterministic and synchronous dynamics. For each *F_i_*, the interactions with all its regulators is based on the biological knowledge described on sections 2.1 and 2.2. The list of all regulatory functions is presented in the supplementary material. A concrete example of the construction of one regulatory function is presented in the Appendix A of the supplementary material.

The *state* of the network *X(t)* at time *t*, is defined as the value that each variable *x_i_* takes at time *t*. That is, *X (t)* = *x_1_(t), x_2_(t), …, x_N_ (t)*. Starting from any initial state of the network *X(0) = x_1_(0), x_2_(0), …, x_N_(0)*, the synchronous and deterministic application of equation (1) results in one and only one successor state for *X(t)*, which is *X(t + 1)*. Iterative application of equation (1) to calculate successor states drives the network through a series of changes that end in a so-called *attractor*, which can potentially be either a *fixed point* if the state does not change in time, or a *cyclic attractor* if a set of states are periodically visited in order. There can be many attractors for the same network. In our model there are no fixed points and only cyclic attractors are present. All the states that under the dynamics end in a specific attractor are its *basin of attraction*. Due to the deterministic nature of the dynamics, a state can only belong to one basin of attraction. This means that the initial state of the network completely determines the attractor that the network will eventually reach. States, attractors, basins of attraction and transitions between states can be represented by a global *state transition graph*. In this graph, each node represents a state and the directed link connecting two states denotes the state transition from one to the other in a single time step. Figure (3) exemplifies a component of the state transition graph for one attractor and its basin of attraction.

**Figure 3:**
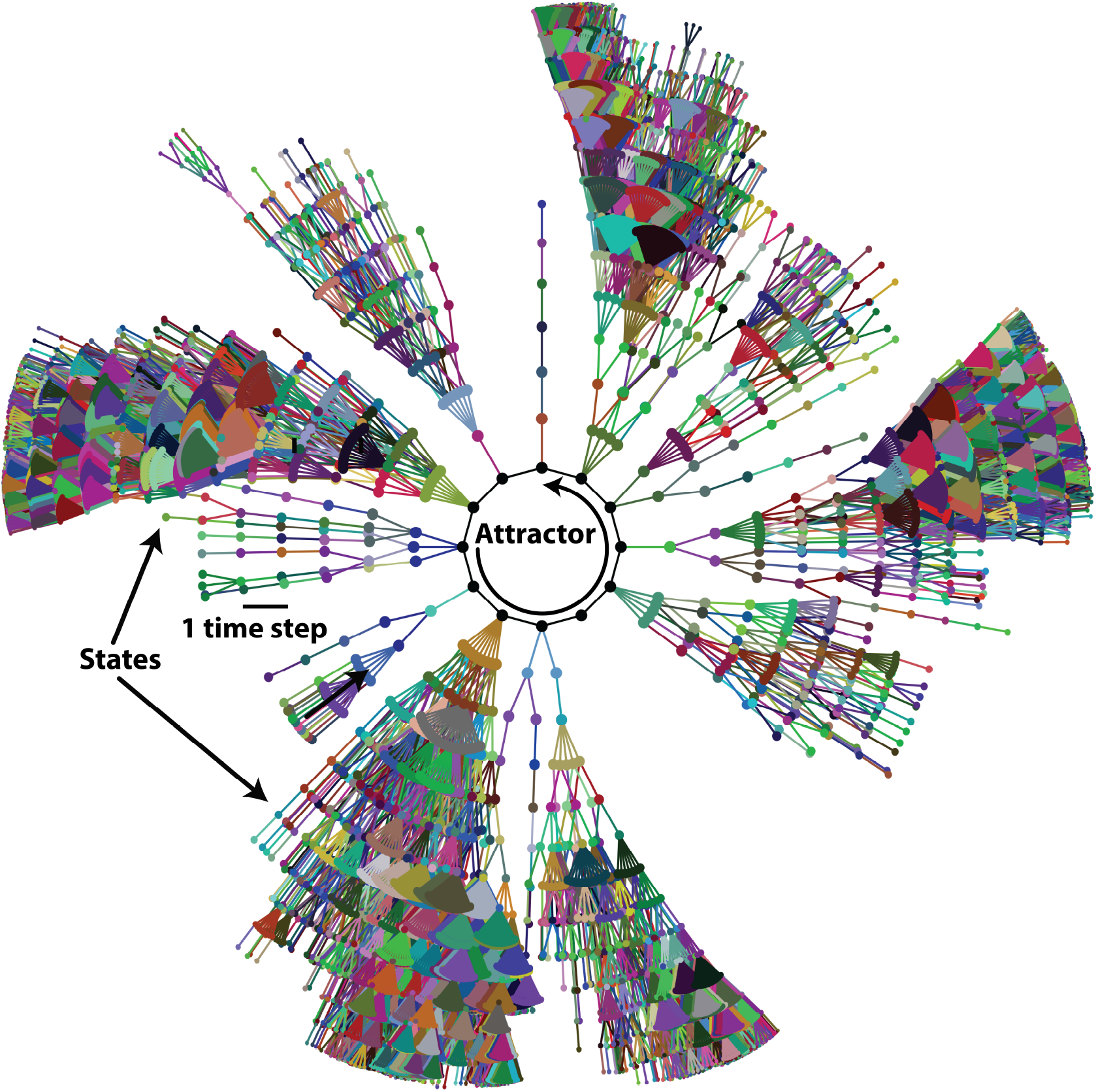
Basin of attraction of one attractor of the AR network. Network states are symbolized by dots and links between dots represent the single-step state transitions. The network eventually reaches the attractor, symbolized by black dots at the center, connected with solid black lines. For clarity, only states at thirteen or less time steps away from the attractor are shown.

It has been shown that different attractors can represent distinct functional behaviors of the system. In genetic regulatory networks the attractors represent patterns of genetic activity that produce different phenotypes [60, 61]. In our case, each state *X(t)* represents the biochemical configuration *x_1_,x_2_, … ,x_N_* of one spermatozoon at time t and attractors represent the different activity patterns a spermatozoon can reach.

## 3 Results

### 3.1 Attractor space partitions into three functional subpopulations

Imagine a population of sperm, each running internally its own regulatory network. We start each network in the population from a random initial condition and let the dynamics run. Different networks will end up in different attractors depending on the initial condition they started from. Thus, although all the networks in the population are identical, the final configuration of the population will be heterogeneous as different networks will end up in different attractors.

To consolidate the construction of the discrete regulatory network model of the human sperm AR, we first characterized and classified the network attractors and their corresponding basins of attraction.

Pg released by the cumulus cells under physiological conditions [4, 59] activates CatSper and therefore increases [Ca^2+^]_*i*_, though other responses have been invoked in the literature [88], promoting the AR. For this reason, we calculated the network attractors in the presence and absence of Pg, by setting the corresponding input node to the active or inactive state. In each case, we identified the attractors in which the reporter node Fusion was active (F=1), that is, attractors that represent sperm that have completed the AR. Figure (4.A) shows the distribution of the 69 different attractors under these conditions. As exemplified in Figure (4.C), these attractors fall into three main *functional classes* in terms of their biological interpretation:

- *Spontaneous*, where the reporter node Fusion activates without activity of the Pg node. Networks in these attractors represent sperm that undergo AR spontaneously without the need of any external stimuli.
- *Inducible*, where activation of the reporter node Fusion requires activation of the Pg node. Networks in these attractors represent sperm that require addition of Pg to undergo AR.
- *Negative*, where the reporter node Fusion never activates, even when the Pg node is active. Networks in these attractors represent sperm that do not react even in the presence of Pg.

**Figure 4:**
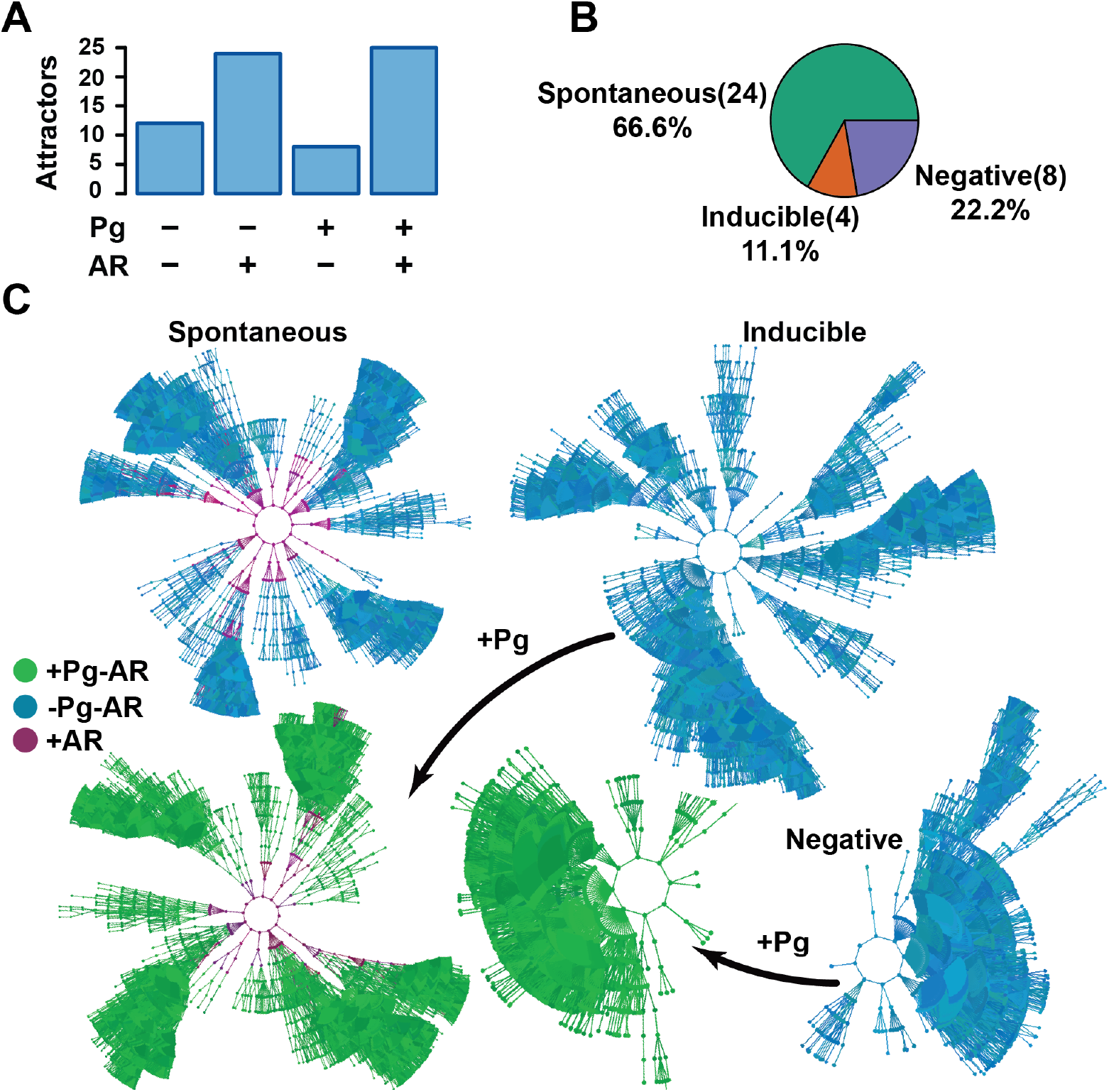
Attractor landscape partitions into three functional groups: Spontaneous, Inducible and Negative. **A)** Total possible attractors classified by their AR activity (F=1) in presence or absence of Pg. **B)** Distribution of attractors in absence of Pg in terms of their functional group. **C)** Diagram of the classification criteria of the attractor space in three functional groups. Colors represent the activity of the Pg node and the reporter node Fusion; blue dots correspond to states where Pg is inactive (Pg=0), whereas green dots indicate active Pg (Pg=1). Purple dots represent states in which membrane fusion has been reached (F=1). Attractor classification follows the next rule: Spontaneous attractors are those where the reporter node Fusion activates when Pg node is inactive, Inducible attractors are those where the reporter node Fusion does not activate unless the Pg node is activate and in Negative attractors the reporter node Fusion is inactive irrespective of the value of the Pg node.

Figure (4.B) shows the distribution of attractors in each of these functional groups according to our classification criteria.

### 3.2 Biochemical heterogeneity and model validation

Human sperm seem to be heterogeneous in their capacity to undergo AR *in vitro*. About 15-20% of the capacitated sperm spontaneously acrosome react [69] and 20-30% of the cells undergo AR in response to Pg [88]. However, the relation between the biochemical heterogeneity in a sperm population and the proportion of cells undergoing spontaneous and Pg induced AR has not been established.

As in different discrete models reported in the literature[1, 29, 30, 31, 61], we used a uniformly distributed random set of network initial states to evaluate the possible fates of the network. States representing cells that have already undergone membrane fusion or that have a fusion machinery already activated, were excluded from the analysis, since these nodes are mostly insensitive to any stimuli and are biologically unreasonable. Therefore we restricted the set of initial states to those where the values of the reporter node Fusion, Swell and SNARE nodes were 0. The values for all the other nodes were assigned randomly.

As shown by Figure (5), the proportion of initial states reaching Inducible, Spontaneous and Negative attractors departed from the experimental observed proportions of cells undergoing Pg-induced AR, spontaneous AR and no AR at all. This discrepancy indicates that it is unrealistic to assume a uniform equiprobable distribution of initial states, and that the set of initial states, interpreted as the heterogeneity in the initial sperm biochemical states, must have a non-trivial distribution. Nevertheless, the results indicate that the biochemical heterogeneity is closely related with the proportion of cells displaying AR, either spontaneous or induced.

**Figure 5:**
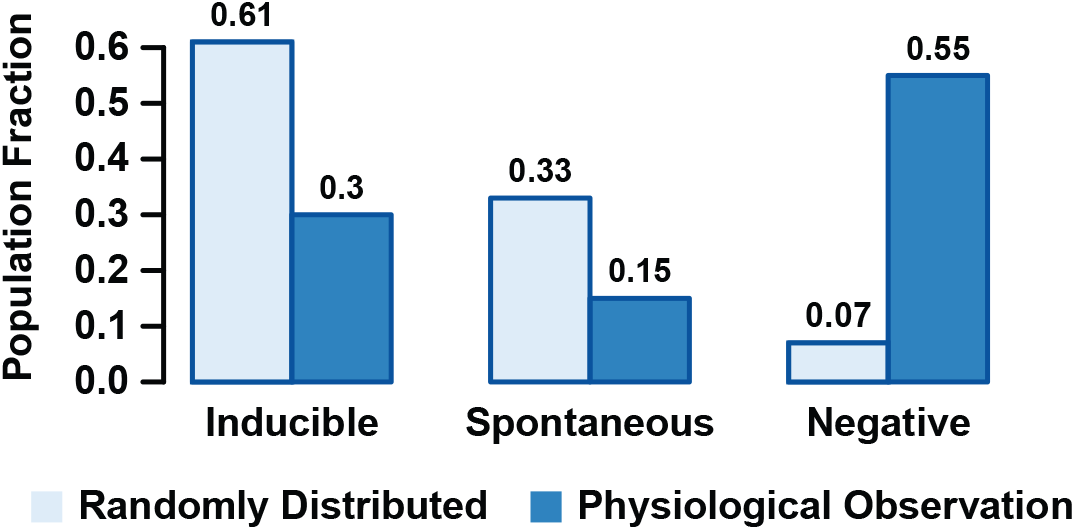
Fraction of states randomly selected based on an equiprobable random distribution that are contained in the basins of attraction of Inducible, Spontaneous and Negative attractors and comparison with the experimental observed fraction of sperms displaying induced, Spontaneous or Negative AR.

To gain insight into the more realistic heterogeneity pattern, we selected the set of initial states by randomly drawing states from the basins of attraction of the Spontaneous, Inducible and Negative attractors, based on probabilities that matched the observed percentages of 15%, 30% and 65%, respectively, and imposing the aforementioned restrictions on the fusion machinery nodes. Notice that the initial state selection procedure cancels any effect of the relative sizes of the basins of attractions and that by construction, the population of networks starting from this set of initial states will reproduce the observed proportion of spontaneous, inducible and negative sub-populations. We used this set to complete our validation of the construction of the model as well as to investigate the putative common features and differences between those initial states. For that purpose, we explored first the activation probability *P(x)* of every node *x* in the network. That is:

- For Boolean nodes, we calculated the probability that a node *x* is in active state *P(x) = P(x = 1)* within the set of initial states.
- For the non Boolean nodes pH_*i*_, CatSper, [Ca^2+^]_*i*_ and Em, *P(x) = P(x > M_x_/2)*, where *M_x_* is the maximum value that the node *x* can take.

Figure (6) shows the activation probability of each node in the network (excluding nodes that represent external medium components). To have good statistics, we sampled 2 × 10^7^ different initial states constructed as described above, which kept the standard deviation of the activation probability below 0.001. The nodes can be separated into two main groups: those whose activation probability is between 0.4 and 0.6, not biased to activation or inactivation, and those that have a lesser than 0.4 or larger than 0.6 activation probability, therefore having a strong bias towards being either activated or inactivated. In agreement with the literature [86], nodes downstream of the cAMP signaling cascade (RAP, Rab3A, cAMP, sAC) as well as IP_3_R Ca^2+^ channels were predominantly inactive. On the other hand, IKsper displayed a tendency to be active, which would be required to maintain the membrane hyperpolarized [12]. In addition, the acrosomal Ca^2+^ ATP-ase (ACA) and H^+^ channel Hv show high biases towards activation. ACA helps to maintain high concentrations of Ca^2+^ inside the acrosome, required for AR, while Hv increases pH_*i*_, an event related to CatSper activation and also necessary for AR [22, 53, 63]. Figure (6) also shows the activation probabilities of the nodes in the selected set of initial states (blue bars), in an equiprobable random set of initial states (dashed orange line) and in all the states composing the basins of attractions of all the attractors of the system (purple line). The node activation pattern of the selected initial states diverges from the random initial states pattern in most of the nodes, especially on those nodes downstream of the cAMP signaling cascade, nodes related to Ca^2+^ mobilization as IP_3_R_*a*_, PLC_*δ–ϵ*_, ACA and nodes linked to pH_*i*_ and Em regulation as Hv and IKsper respectively. Although subtle, the difference between the node activation patterns of the selected initial states and the states in the entire basins of attraction is noticeable, where the bigger differences can be appreciated in the nodes related to cAMP and [Ca^2+^]_*a*_ signaling cascades, that show a lower activation probability in the selected set of initial states than in the random set of initial states. This is likely because in the Spontaneous and Inducible attractors, the activation of these nodes favors the activation of the reporter node Fusion. Consequently, the states where cAMP and [Ca^2+^]_*a*_ regulation related nodes are active, are congregated around the attractor. This is the reason that the entire set of states in the basin of attraction considers more states with activation of these nodes than the selected set of initial states, explaining activation probability pattern differences.

**Figure 6:**
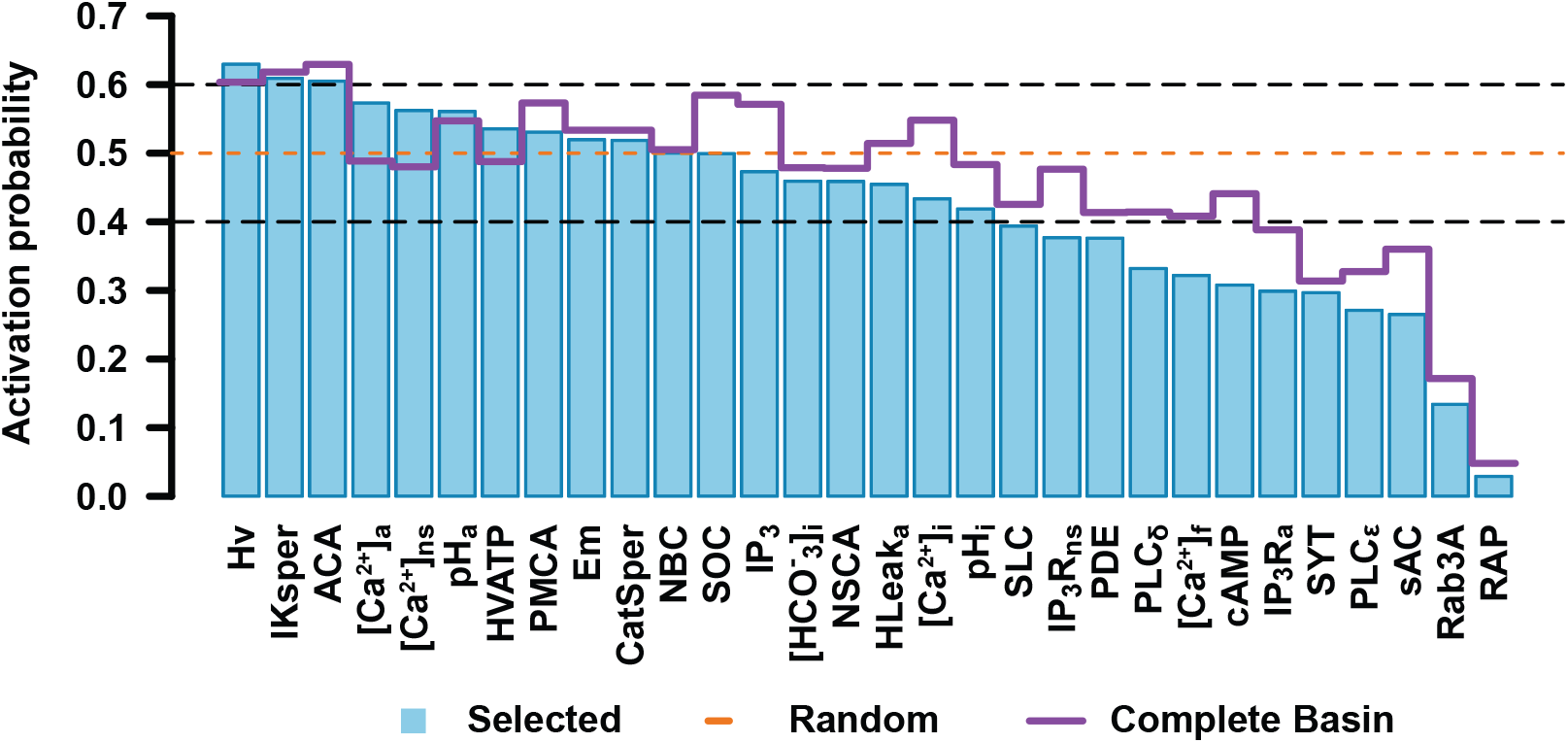
Activation probability of network nodes in the set of initial states. Bar heights for each node represent the probability that the node is in the active state. The orange dashed line points out the activation probability of a completely random set of initial states. The purple straight line represents the activation probability of each node on the entire set of states conforming the basins of attraction for all the attractors. Nodes whose activation probability is between 0.4 and 0.6 are enclosed in black dashed lines. Error bars are not shown since in each case standard deviation is below 0.001.

For non-binary nodes, Figure (7) shows the probability that a node takes a specific value in the set of initial states. As can be seen, only a small population of initial state take the most alkaline pH_*i*_, while most initial states take more acidic values. This agrees with the notion that cytosolic alkalinization is required for AR [22, 33, 53]. Most of the initial states in the population belong to the Negative attractor group that will not lead to either spontaneous or induced AR. Also, only small fractions of the set of initial states show the open state (1) for CatSper and the high [Ca^2+^]_*i*_ levels, agreeing with the idea that CatSper activation is important to increase [Ca^2+^]_*i*_ and AR [4, 64]. No significant differences can be appreciated among the Em values, although this could be compensated by the activity of IKsper shown in Figure 6 to display a strong tendency of Em towards hyperpolarized values in the Inducible attractors (Figure 8). These results suggest that the proportion of a sperm population displaying spontaneous, Pg induced AR or no AR, is related to the sperm biochemical state heterogeneity. Furthermore, this heterogeneity is not arbitrary and there are cell elements more robust to perturbations than others.

**Figure 7:**
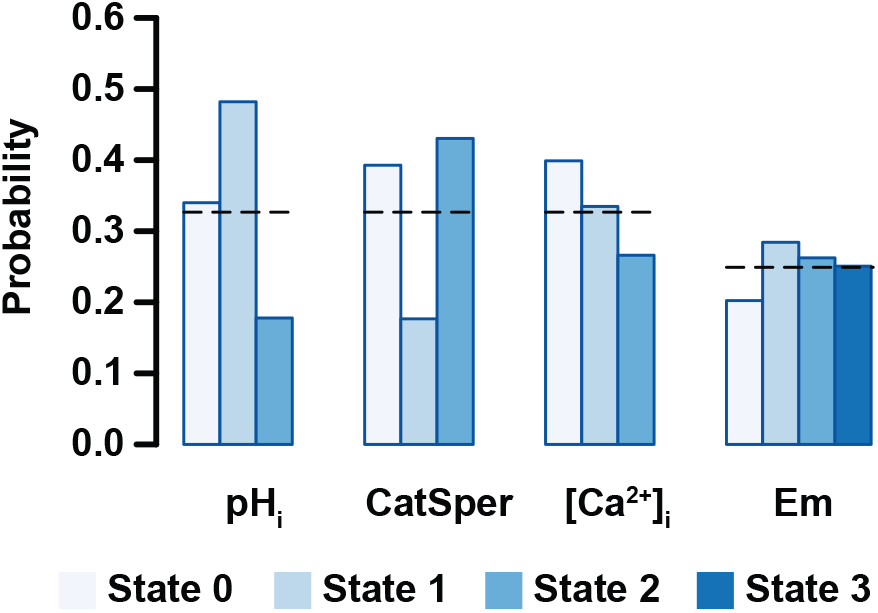
Probability that one node takes a specific value in the set of initial states for non binary nodes. Bar lengths show the average probability calculated over 2 × 10^7^ initial states. Black dashed lines indicate the probability on a completely random set of initial states. Biological interpretation for each node is as follows: pH_*i*_ (acidic 0, mildly alkaline 1, fully alkaline 2); CatSper (closed 0, open 1, inactivated 2); [Ca^2+^]_*i*_ (basal 0, activator 1, inhibitor 2); Em (hyperpolarized 0, equilibrium 1, mildly depolarized 2, and fully depolarized 3). Error bars are not shown since standard deviation is below 0.001.

As mentioned above, to further validate the model and to better characterise the selected set of initial states, we compared the results of the model with a set of 26 different experimental observations reported in the literature and described in supplementary Table (2). For each experimental observation, we constructed a set of networks starting from the aforementioned set of initial states. On each network, we calculated dynamics by successive application of equation 1 until it reached its corresponding attractor. Depending on each experiment, we simulated the effect of specific stimuli by solving the dynamics until the attractor was attained in a scenario where the stimulus-associated node is inactive or active.

We calculated the average trajectory of all the networks with and without the stimulus and compared these responses with the ones reported in the experiments. We focused our attention on the behavior of [Ca^2+^]_*i*_, pH_*i*_, Em nodes and the reporter node Fusion given their relevance for the AR. Table (2) displays the results of these comparisons, showing that our model is in agreement with 92% of the experimental results obtained by different research groups.

**Table 2:**
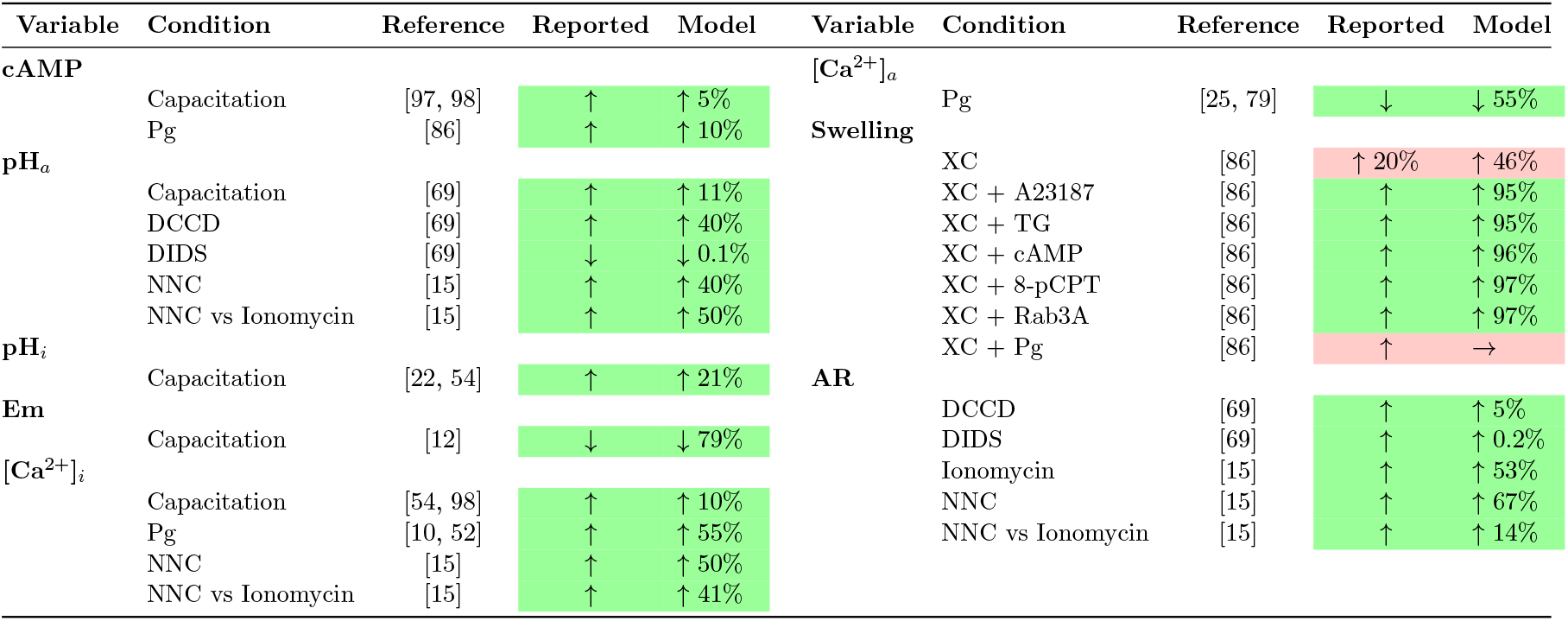
Comparison of the 26 experimental conditions tested in the AR discrete network with reported observations. The model has a success rate of 92%. Values presented in the network observations are relative to maximal activity of the respective node. References from which results were derived are indicated. Percentages are given considering maximum AR under a specific condition as 100% after examining the selected initial states. Arrows represent an increase (↑), a decrease (↓) or no change (→) in the corresponding variable for the reported experiment and the model. Rows in green and red indicate agreement and disagreement respectively, between the model and the reported experimental results. All the experiments were performed in different capacitating mediums (see references). DCCD, N,N’-dicyclohexylcarbodiimide; DIDS, 4,4’-diisothiocyanato-stilbene-2,2’-dislufonic acid; NNC, NNC55-0396; XC, Xestospongin C; TG, Thapsigargin; 8-pCPT, 8-pCPT-20-O-Me-cAMP.

### 3.3 Spontaneous, Inducible and Negative attractors show large scale differences on the network biochemical state

Once having validated our network model we proceeded to examine its prediction capacity. To this end we further analyzed the characteristics that define the functional classification of the attractors into the Spontaneous, Inducible and Negative AR. We calculated the activation probability of all the nodes for each functional group within the set of initial states previously constructed. Figure (8.A) shows the nodes where the activation probability is significantly different among the three functional groups. The biggest differences are shown in the Inducible class, where activity of nodes related to [Ca^2+^]_*i*_ and [Ca^2+^]_*a*_ have the most significant change in the activation probability compared with the other two classes. Although differences between Negative and Spontaneous attractors are subtle, it can be seen that there are considerable changes in the activation probability for sAC, the acrosome H^+^ outward current (HLeak), pH_*i*_, the acrosomal V-ATPase and Em. Notably, this figure shows that, in the set of initial conditions, the nodes related to pH_*i*_ and pH_*a*_ regulation display different activation probabilities among the functional classes.

**Figure 8:**
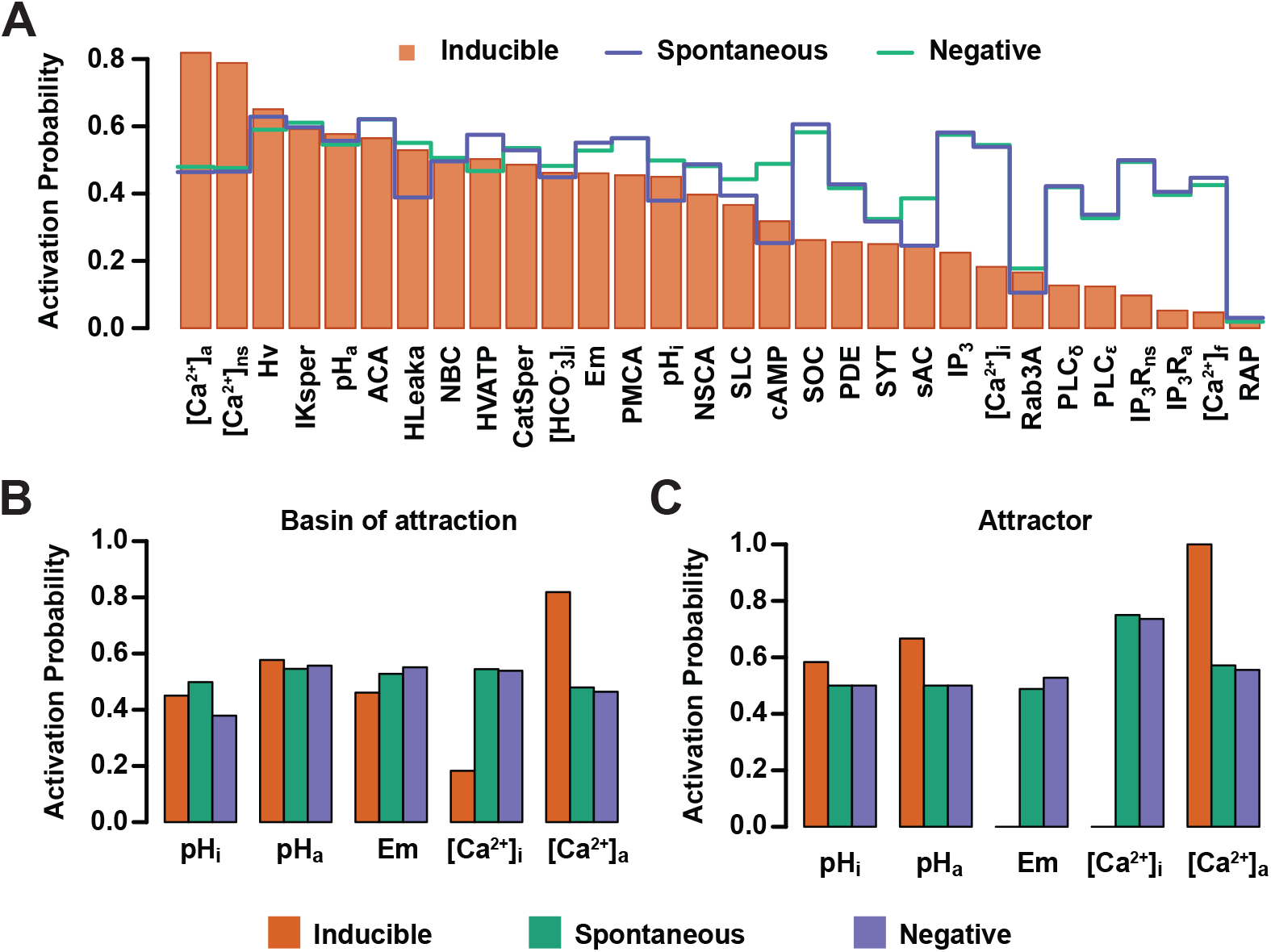
Activation probability of network nodes in Inducible, Spontaneous and Negative attractors. **A)** Bars represent the activation probability of the network nodes, for states in the Inducible attractors. Blue and green lines correspond to the activation probability of the network nodes in the Spontaneous and Negative attractors, respectively. **B-C)** Activation probability of pH_*i*_, pH_*a*_, Em. [Ca^2+^]_*i*_ and [Ca^2+^]_*a*_ in the basins of attraction **B)** or attractors **C)** in the three functional classes.

We focused on the pH_*i*_, pH_*a*_, Em, [Ca^2+^]_*i*_ and [Ca^2+^]_*a*_ nodes given their relevance for AR. Figure (8.B) shows there are significant differences in the activation probability of these nodes between functional classes. As expected, the pH_*i*_ node displays a higher activation probability in the Spontaneous than in the Inducible attractors, being the least likely in the Negative class. No significant differences can be appreciated regarding the pH_*a*_ node activation probability among the three classes. Remarkably, the Em and [Ca^2+^]_*i*_ nodes reach their lowest activation probabilities in the Inducible class, indicating that increases in Em activity (depolarization) and in [Ca^2+^]_*i*_ activity (increased concentration) often lead the system to attractors that display spontaneous fusion activity or no fusion activity at all. The [Ca^2+^]_*a*_ node shows its highest value also in the Inducible class, indicating that low [Ca^2+^]_*a*_ node activity drives the network to the Spontaneous or Negative attractors. This behavior is mostly preserved when we apply the same procedure, not to the entire basin of attraction, but to the attractor itself, calculating the activation probability of the nodes within the attractor’s states. As Figure (8.C) shows, differences in this case are more evident between classes; the pH_*i*_ and pH_*a*_ nodes display their highest activity levels in the Inducible attractors. We found that in all the network states of the attractors in this class, Em is at its lowest value (Em=0), representing hyperpolarization, which means that whenever an attractor state contains a value for Em different than 0, the dynamics drive the network either to the Spontaneous or Negative attractors. Similarly, the [Ca^2+^]_*i*_ node always takes the basal value 0 in attractors of the Inducible class. This does not oppose established evidence that [Ca^2+^]_*i*_ rises during capacitation but indicates it must remain at basal levels while awaiting for a stimulus such as Pg to trigger AR, otherwise, an increase in the activity of the [Ca^2+^]_*i*_ node drives the network to a Spontaneous or Negative attractor. As expected, the [Ca^2+^]_*a*_ node activity tends to be lower within the Spontaneous attractors and it reaches its highest value in the Inducible attractors.

These results are consistent with the notion that AR reaction is promoted by increases in [Ca^2+^]_*i*_ as well as membrane depolarization, but they also suggest the existence of hyperpolarized sperm with elevated [Ca^2+^]_*i*_ that do not react. The activation probability of the [Ca^2+^]_*a*_ node in the Inducible class is consistent with the idea that acrosomal stability requires high Ca^2+^ levels to prepare sperm for the Pg stimulus, and contributing to maintain low [Ca^2+^]_*i*_. Also, the pH_*i*_ node activation probability pattern displayed in the functional classes agrees with the notion that a cytosol alkalinization enables spontaneous as well as Pg-induced AR, but the model predicts a more elevated pH_*i*_ in cells that can respond to Pg than in cells that undergo AR spontaneously.

### 3.4 Elevation of pH_*a*_ defines network capacity for AR

To further investigate the role of pH_*a*_ during acrosomal exocytosis, we analyzed how the attractor landscape changes as a consequence of acidic (pH_*a*_=5.5) or alkaline (pH_*a*_=6.7) conditions by fixing the pH_*a*_ node value to 0 or 1, respectively. We noticed first, that fixing the pH_*a*_ node drastically reduced the number of total attractors, as shown in Figure (9.A). Without determining a particular value for pH_*a*_, and setting Pg=0, which in our model represents absence of Pg, the dynamics generate a total of 36 attractors that are reduced to 8 and 10 in conditions representing low (0-acidic) and high (1-alkaline) pH_*a*_, respectively. Remarkably, the extent of this contraction does not follow from the fact that fixing the value of pH_*a*_ reduces by half the total number of possible network states. This change implies that fixing pH_*a*_ limits the attractor landscape and therefore reduces the possible different stable behaviors that the network can reach from any of the available initial states.

**Figure 9:**
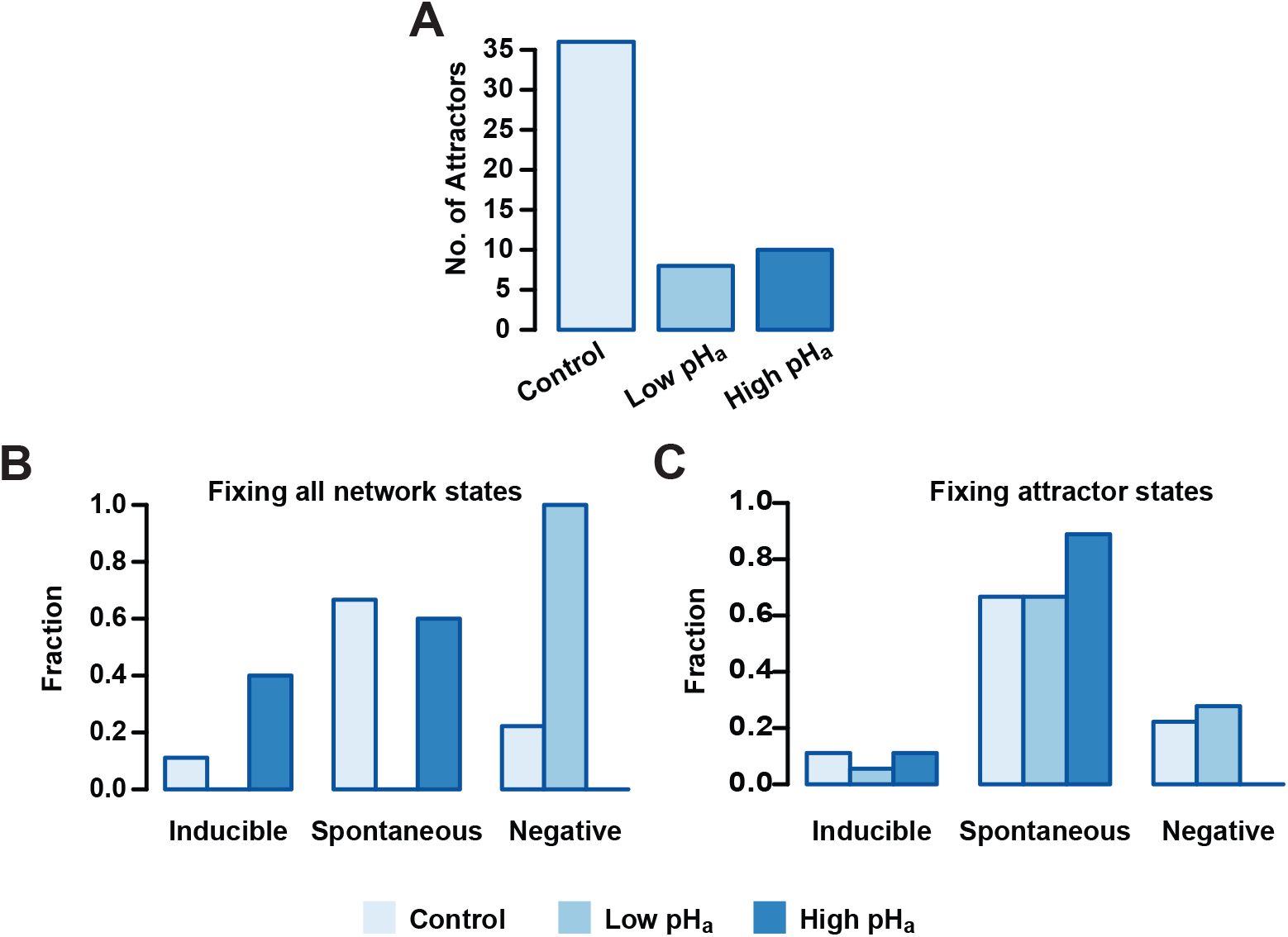
pH_*a*_ blocking effect on Inducible, Spontaneous and Negative basins of attraction. **A)** Number of total attractors when the pH_*a*_ node is fixed as acidic (0) or alkaline (1) on the entire state space. **B)** Distribution of attractors among functional classes fixing pH_*a*_ as in A. Setting pH_*a*_=0 results in the deletion of Inducible and Spontaneous attractors, whereas fixing pH_*a*_=1 leads to deletion of the Negative class. **C)** Attractors distributed among the different functional classes where pH_*a*_ is fixed only in the attractor states. Fixing pH_*a*_=1 on the sequence of Negative attractors transforms them in Spontaneous attractors.

To better understand the repercussions of this landscape contraction, we re-classified the resulting attractor landscape according to the functional classes and we calculated the fraction of the total attractors that resides in each one. Figure (9.B) shows that for pH_*a*_ = 0, no attractor displays activation of the reporter node Fusion on any of its states, and the entire attractor landscape falls into the Negative AR class. On the other hand, the Negative class vanishes when pH_*a*_ = 1 and all attractors in the landscape display activation of the reporter node Fusion, either in the Spontaneous or Inducible classes. Noticeably, only Inducible attractors increase with respect to the control group in this condition, suggesting that only the induced AR is promoted by acrosomal alkalinization.

Constraining the pH_*a*_ node to a fixed value on the entire state space is a major perturbation of the system since it trims the space of available states and reduces the total number of attractors. Considering this, we also analyzed a less drastic perturbation: instead of fixing pH_*a*_ on all possible states, we fixed it only on the states conforming to the original 36 attractors. This would mean altering pH_*a*_ once the cell reached an attractor after being capacitated. As shown in Figure (9.C), this perturbation has less drastic effects on the functional distribution of the attractors, but still, when pH_*a*_ = 0, the number of Inducible attractors decreases and the number of Negative attractors increases, while the number of Spontaneous attractors remains the same. This means that the dynamics under this condition reduces the capacity of the network to activate the reporter node Fusion under Pg activation. In contrast, the Negative class disappears when pH_*a*_ = 1 and the number of Spontaneous attractors increases while the Inducible class remains intact. These findings indicate that a pH_*a*_ increase can by itself trigger AR in a group of cells, but also, that it increases the number of cells that make AR induced by Pg.

### 3.5 Calcium and pH transitory perturbations promote functional changes with the highest probability

At this point we should recall that the dynamics in our model, described by equation (1) is deterministic and synchronously updated. This means that each initial state leads to one and only one attractor. As a further step in understanding how robust the network is to small transitory changes in pH_*a*_, as well as how it compares to the effect of the same kind of changes in other nodes, we implemented a series of small node-specific perturbations and we counted the fraction of those perturbations that promoted not only a change in the residing basin of attraction but also in the functional class of the resulting attractor. To this purpose we took a set of 10^5^ random states in the basin of attraction of each attractor and changed the value of one single node, the same for all states. In the case of binary nodes we simply inverted the value for its complement; for non-binary nodes we randomly took a different value among the set of possible node values. In the particular case of the Spontaneous class, given the irreversibility of membrane fusion, we took our sample only from those states that did not translate into a final AR stage, that is, we only selected those conditions where the nodes Swelling, SNARE’s, SYT and the reporter node Fusion had a value of 0.

The results shown in Figure (10) present in order the nodes where the perturbations had the highest rates of change in the attractor type for the different functional classes. As shown in panel A, the nodes that made the Inducible attractors more susceptible to perturbation were those related to the regulation of the [Ca^2+^]_*i*_ and [Ca^2+^]_*a*_. Remarkably, pH_*a*_ node perturbations did not promote a significant number of functional changes. Also, most of the changes promoted were to the Spontaneous class. Panels B and C show that in the cases of Spontaneous and Negative attractors, the nodes most susceptible to perturbation were mainly pH_*a*_ and those related to pH_*i*_ and pH_*a*_ regulation. Panel B illustrates that there is only a small amount of changes produced by perturbations in the Spontaneous attractors that can lead the network to the Negative class. Noticeably, most of the changes in the Negative attractors were directed towards the Spontaneous attractors as shown in panel C. This result predicts that changes in pH_*a*_ can promote AR, at least partially, without any other stimuli and that most of the sperm reacted in this way come from a group of sperm that would have not displayed Spontaneous or Pg induced AR. The results presented in this section indicate that not only sustained changes promote AR but also transitory stimuli, mostly in Ca^2+^ and pH in cytosol and acrosome are important as inducers of the AR. We address this topic further in the discussion.

**Figure 10:**
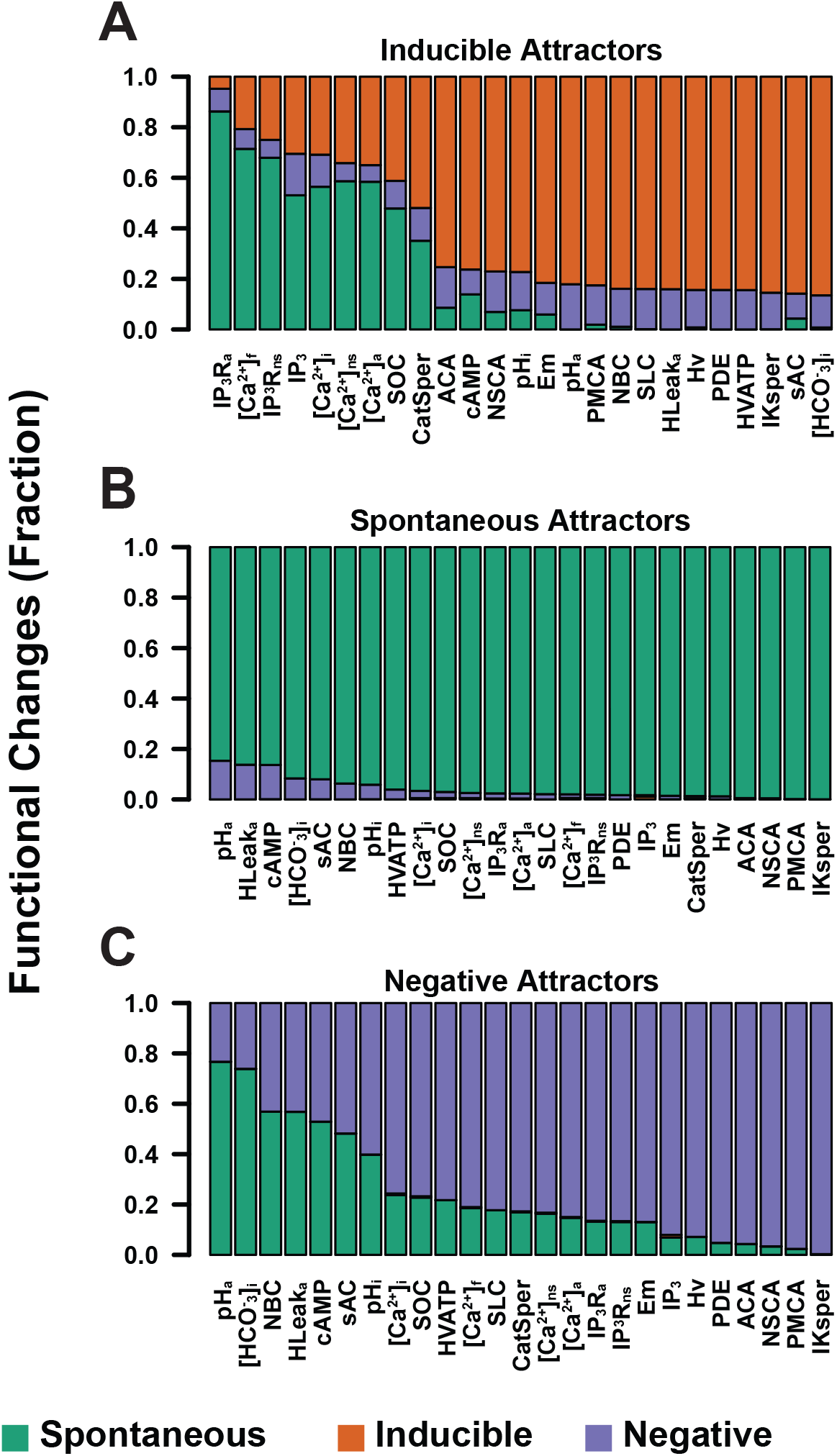
Fraction of node-specific perturbations that promoted functional changes on Inducible, Spontaneous and Negative attractors. Nodes involved directly in [Ca^2+^]_*i*_ and [Ca^2+^]_*a*_ regulation show high sensitivity on the Inducible group, whereas Spontaneous and Negative attractors show higher sensitivity to perturbations to nodes related to pH_*a*_ and pH_*i*_. The three panels show a minimal change into Inducible attractors from Negative and Spontaneous groups.

## 4 Discussion

In this work, we developed a discrete, synchronous and deterministic regulatory network for the acrosome reaction in human sperm. The set of biochemical pathways describing the processes that culminate in the fusion of the outer acrosomal membrane and the plasma membrane are far from clear and sometimes the existing evidence is controversial. However, the network presented here, takes into account the most recent experimental evidence reported in the literature regarding the main intracellular components and their interactions involving this complex exocytotic process, with the hope that it may promote its better understanding. Further integration of novel information and data into the model could help to clarify the signaling cascades participating in the human sperm acrosome reaction.

We calculated the attractor landscape and identified the states that belong to their corresponding *basins of attraction*. We concluded that this network has only cyclic attractors representing stable biochemical oscillations, there are no fixed points. We noticed that these attractors are modified by the addition of Pg to the extracellular medium, and that these differences are consistent with the response displayed experimentally by a sperm population under the same stimulus. Taking into account this finding, we partitioned the attractor landscape in Spontaneous, Inducible and Negative attractors based on their functional characteristics.

As is common in the logical dynamics literature [1, 29, 30, 31, 61], we addressed the probabilities reaching different attractors by taking random and uniformly distributed initial states from the state transitions graph. This approach led us to to conclude that the proportions of network populations reaching spontaneous and Pg-induced AR are inconsistent with the proportions observed experimentally [69, 88]. We explored the characteristics of different sets of initial states that were more consistent with proportions of AR experimentally reported under physiological conditions. We determined that the activation probability of each node in this set of initial states agrees with the biological behavior reported in the literature. This suggested that not all of the possible network biochemical states are reachable under physiological conditions. Moreover, the number of attractors displayed by the network and the size of their basins of attraction is not physiologically relevant. With our study we were able to delimit the set of network states that should be considered as potential states characteristic of capacitated sperm.

Furthermore, our results showed that, within the set of initial states, a group of nodes displayed an activation probability close to 0.5, meaning that it is equally probable that they take any of their possible values, while for other nodes the activation probability has a strong bias towards either basal levels (0) or increased activity (> 0). In all the sets of initial states analyzed, these activation patterns are required in order to reproduce the experimental proportions of spontaneous AR, Pg induced AR and no AR at all. This suggests that there is a relation between the structure and characteristics of the heterogeneity in the biochemical states in a sperm population and how this population is partitioned to display the observed proportions. As stated before, our model considers events between later stages of capacitation and membrane fusion and it is not suitable to investigate the origin of population heterogeneity, how it changes during the different stages in sperm development and maturation and the order of events that finally promote these activity patterns. To address these matters, our model can be expanded to consider the events that take place during early capacitation stages or even spermatogenesis. This would allow us to evaluate the structure of heterogeneity at different moments during the sperm maturation processes.

We showed that the model can qualitatively reproduce experimental observations reported in the literature. In this sense, our model has an overall success rate of 92% in reproducing 24 of 26 experimental results describing different behaviors of a sperm population in response to diverse conditions and stimuli.

An experimental observation our model was unable to reproduce is the finding that Pg increases acrosomal swelling in a population of human sperm in the presence of xestopongine C (XC), an IP_3_R blocker [86]. In our model, Pg promotes only the activity of CatSper in the sperm flagellum [4, 59]. We hypothesized that the local increase in flagellar [Ca^2+^]_*i*_ and its diffusion to the head is insufficient to promote the activation of the signaling cascade that triggers the fusion machinery, which involves a Ca^2+^ dependent cAMP increase and a Ca^2+^ dependent IP_3_R channel. Instead, we envisage that the local Ca^2+^ wave generated in the flagellum is re-transmitted by IP_3_R located in the neck stores and that this re-transmission, being closer to the head, provides the necessary Ca^2+^ to activate AR. However, in presence of XC this would not be possible, as this inhibitor would stop re-transmission from the neck stores and thus, the flagellar Ca^2+^ signal would not reach the head, preventing acrosome swelling and AR to occur. This result could suggest the participation of a different channel involved in the retransmition of the Ca^2+^ signal, located in the neck stores. For example, there is evidence suggesting that the Ryanodine receptors (RyRs), located in the redundant nuclear envelope in human sperm, are involved in the generation of intracellular spontaneous and Pg induced [Ca^2+^]_*i*_ oscillations[57]. However, the role and expression of RyRs in human sperm functioning has been controversial [6, 39, 57, 91]. The discrepancy between our model and this particular experiment could also suggest that Pg promotes a [Ca^2+^]_*i*_ increase in the head or activates AR by a different pathway. It has been shown that Pg promotes [Ca^2+^]_*i*_ increases through CatSper in humans [52, 89], nevertheless this hormone does not activate CatSper in mice [56, 74], although it can still promote AR [66, 67]. However, the alternate signaling path triggered by Pg in the head and its relation to the AR is unknown.

Next, we calculated the probability that each node in the network is active in the Spontaneous, Inducible and Negative attractors. The differences in the activation probability give us insight on the differences in the biochemical state of sperm and their capacity to reach AR. We found that the attractors in the Inducible group have notable differences, especially in the Em, [Ca^2+^]_*i*_, [Ca^2+^]_*a*_, pH_*a*_ and pH_*i*_ nodes, which is consistent with the literature.

We tested the behavior of the attractor landscape distribution among functional classes to identify how attractors are affected by changes in the state of the pH_*a*_ node. We found that the total number of attractors is drastically reduced when low and high pH_*a*_ conditions are simulated and applied to the entire space of possible network states. As this is a drastic maneuver, we also tested fixing the value of the pH_*a*_ node only in the states that conform the attractor to determine how attractors distributed among the functional classes, while preserving the total number of attractors. In both cases we found that the conditions where pH_*a*_ = 0 increase the number of Negative attractors, preventing the network from activating the reporter node Fusion. This means that acidic pH_*a*_ conditions would partially prevent AR. Remarkably, when pH_*a*_ = 1, the Negative attractors disappeared and were transformed into Inducible or Spontaneous attractors. These results suggest that for a fraction of the sperm population, acrosomal alkalinization acts as a necessary step for AR induction. In addition, they indicate that a pH_*a*_ rise is partially promoting AR by itself in a group of sperm. This model prediction is supported by published work from our group [15].

We then applied a series of small and transitory perturbations to the states in the basins of attraction of the different functional classes of attractors, to identify the network elements that are more sensitive to perturbations. We explored if the temporary alteration of a single node could change the trajectory of the network towards an attractor of a different functional class. As it is well known that AR is effectively induced by increasing [Ca^2+^]_*i*_ in capacitated sperm[4, 64], we found that nodes related to [Ca^2+^]_*i*_ and [Ca^2+^]_*a*_ regulation show the highest sensitivity to perturbations and most changes in those nodes promote AR. This findings are consistent with experimental results. Noticeably, we found that pH_*a*_ is the most sensitive node for the Spontaneous and Negative attractors, followed by nodes related to pH_*i*_ and pH_*a*_ regulation. These results suggest that not only long term pH_*a*_ changes induce AR but also, transitory perturbations promote this reaction in a fraction of the sperm population.

Synchronous and deterministic discrete updating are important limitations of this model. Although many elements presented in the network are taken into account to update its state in a synchronized deterministic manner (for example, Em reacts instantly to ionic current changes), there is no evidence indicating this is the case for many other of its elements. Also, the discretization of variables makes it difficult to address some of the problems dealt with here in a more quantitative way. We are in the process of developing a more descriptive model by incorporating stochasticity to non-synchronous and semi-synchronous models.

## Author Contributions

A.A. Conceptualization, Investigation, Methodology, Formal Analysis, Software development, Writing–Review and Editing.

J.C. Conceptualization, Investigation, Methodology, Funding Acquisition, Supervision, Writing–Review and Editing.

G.M-M. Conceptualization, Investigation, Methodology, Funding Acquisition, Supervision, Writing–Review and Editing.

A.D. Conceptualization, Investigation, Methodology, Funding Acquisition, Supervision, Writing–Review and Editing.

## Acknowledgments

We thank Maximino Aldana and Andrea Falcón-Cortés for comments and revision of this paper. A.A and G.M-M thank the hospitality of Jorge Kurchan and the Physics Department of the École Normale Superiéure in Paris (ENS), during an academic stay (A.A) and a sabbatical leave (G.M-M) where part of this research was undertaken. A.A and A.D thank the hospitality of the Instituto Gulbenkian de Ciência, Oeiras, Portugal, during an internship (A.A) and a sabbatical leave (A.D) where part of this investigation was done. We thank Alejandro Aguado-García, Daniel A. Priego-Espinosa, Aurelien Naldí, Denis Thieffry and Claudine Chaouiya for fruitfull discussions. A.A is a doctoral student from the Programa de Doctorado en Ciencias Biomédicas, Universidad Nacional Autónoma de México (UNAM).

## References

[1] E. R. Álvarez-Buylla, Á. Chaos, M. Aldana, M. Benítez, Y. Cortes-Poza, C. Espinosa-Soto, D. A. Hartasánchez, R. B. Lotto, D. Malkin, G. J. Escalera Santos, and P. Padilla-Longoria. Floral morphogenesis: Stochastic explorations of a gene network epigenetic landscape. PLoS ONE, 3(11), 2008.

[2] M. Balbach, M. G. Gervasi, D. Martin-Hidalgo, P. E. Visconti, L. R. Levin, and J. Buck. Metabolic changes in mouse sperm during capacitation. Biology of Reproduction, 1(11), 2020.

[3] M. Balbach, H. Hamzeh, J. F. Jikeli, C. Brenker, C. Schiffer, J. N. Hansen, P. Neugebauer, C. Trötschel, L. Jovine, L. Han, H. M. Florman, U. B. Kaupp, T. Strünker, and D. Wachten. Molecular Mechanism Underlying the Action of Zona-pellucida Glycoproteins on Mouse Sperm. Frontiers in Cell and Developmental Biology, 8, 2020.

[4] E. Baldi, M. Luconi, M. Muratori, S. Marchiani, L. Tamburrino, and G. Forti. Nongenomic activation of spermatozoa by steroid hormones: Facts and fictions. Molecular and Cellular Endocrinology, 308(1-2):39–46, sep 2009.

[5] R. W. Baxendale and L. R. Fraser. Evidence for multiple distinctly localized adenylyl cyclase isoforms in mammalian spermatozoa. Molecular Reproduction and Development, 2003.

[6] K. Bedu-Addo, S. Costello, C. Harper, G. Machado-Oliveira, L. Lefievre, C. Ford, C. Barratt, and S. Publicover. Mobilisation of stored calcium in the neck region of human sperm a mechanism for regulation of flagellar activity. The International Journal of Developmental Biology, 52(5-6):615–626, 2008.

[7] S. A. Belmonte, L. S. Mayorga, and C. N. Tomes. The Molecules of Sperm Exocytosis. Advances in anatomy, embryology, and cell biology, 220:71–92, 2016.

[8] I. Bezprozvanny, J. Watras, and B. E. Ehrlich. Bell-shaped calcium-response curves of Ins(1,4,5)P3- and calcium-gated channels from endoplasmic reticulum of cerebellum. Nature, 351(6329):751–4, jun 1991.

[9] E. Bianchi and G. J. Wright. Sperm Meets Egg: The Genetics of Mammalian Fertilization. Annual Review of Genetics, 50(1):93–111, 2016.

[10] P. F. Blackmore, S. J. Beeben, D. R. Danforthn, and N. Alexanderll. Progesterone and 17wHydroxyprogesterone NOVEL STIMULATORS OF CALCIUM INFLUX IN HUMAN SPERM. The Journal of Biological Chemistry, 265(3):1376–1380, 1990.

[11] M. T. Branham, L. S. Mayorga, and C. N. Tomes. Calcium-induced acrosomal exocytosis requires cAMP acting through a protein kinase A-independent, Epac-mediated pathway. Journal of Biological Chemistry, 281(13):8656–8666, 2006.

[12] J. C. Chá vez, J. L. de la Vega-Beltrá, J. Escoffier, P. E. Visconti, C. L. Treviñ, A. Darszon, L. Salkoff, and C. M. Santi. Ion Permeabilities in Mouse Sperm Reveal an External Trigger for SLO3-Dependent Hyperpolarization. PLoS ONE, 8(4):60578, 2013.

[13] M. R. Chae, S. J. Kang, K. P. Lee, B. R. Choi, H. K. Kim, J. K. Park, C. Y. Kim, and S. W. Lee. Onion (Allium cepa L.) peel extract (OPE) regulates human sperm motility via protein kinase C-mediated activation of the human voltage-gated proton channel. Andrology, 5(5):979–989, 2017.

[14] C. Chaouiya, A. Naldi, and D. Thieffry. Logical modelling of gene regulatory networks with GINsim. Methods in Molecular Biology, 2012.

[15] J. C. Chávez, J. L. De la Vega-Beltrán, O. José, P. Torres, T. Nishigaki, C. L. Treviño, and A. Darszon. Acrosomal alkalization triggers Ca2+ release and acrosome reaction in mammalian spermatozoa. Journal of Cellular Physiology, 2017.

[16] J. C. Chávez, J. J. Ferreira, A. Butler, J. L. De La Vega Beltrán, C. L. Treviño, A. Darszon, L. Salkoff, and C. M. Santi. SLO3 K+ channels control calcium entry through CATSPER channels in sperm. Journal of Biological Chemistry, 289(46):32266–32275, 2014.

[17] Y. Chen, M. J. Cann, T. N. Litvin, V. Lourgenko, M. L. Sinclair, L. R. Levin, and J. Buck. Soluble Adenylyl Cyclase as an Evolutionarily Conserved Bicarbonate Sensor. Science, 289(5479):625–628, jul 2000.

[18] J. Correia, F. Michelangeli, and S. Publicover. Regulation and roles of Ca2+ stores in human sperm. Reproduction, 150(2):R56–R76, 2015.

[19] R. Da Costa, D. Botana, S. Piñero, F. Proverbio, and R. Marín. Cadmium inhibits motility, activities of plasma membrane Ca2+-ATPase and axonemal dynein-ATPase of human spermatozoa. Andrologia, 48(4):464–469, 2016.

[20] J. C. Dan. Studies on the Acrosome. I. Reaction to Egg-Water and Other Stimuli. Biological Bulletin, 103(1):54–66, 1952.

[21] J. C. Dan. Studies on the acrosome. III. Effect of calcium deficiency. The Biological Bulletin, 107(3):335–349, 1954.

[22] A. Darszon, J. J. Acevedo, B. E. Galindo, E. O. Hernández-González, T. Nishigaki, C. L. Treviño, C. Wood, and C. Beltrán. Sperm channel diversity and functional multiplicity, 2006.

[23] A. Darszon, P. Labarca, T. Nishigaki, and F. Espinosa. Ion Channels in Sperm Physiology. Physiological Reviews, 79(2), 1999.

[24] A. Darszon, T. Nishigaki, C. Beltran, and C. L. Treviño. CALCIUM CHANNELS IN THE DEVELOPMENT, MATURATION, AND FUNCTION OF SPERMATOZOA. Physiological Reviews, 91:1305–1355, 2011.

[25] G. De Blas, M. Michaut, C. L. Treviño, C. N. Tomes, R. Yunes, A. Darszon, and L. S. Mayorga. The Intraacrosomal Calcium Pool Plays a Direct Role in Acrosomal Exocytosis. Journal of Biological Chemistry, 277(51):49326–49331, dec 2002.

[26] G. A. De Blas, C. M. Roggero, C. N. Tomes, and L. S. Mayorga. Dynamics of SNARE assembly and disassembly during sperm acrosomal exocytosis. PLoS Biology, 3(10), 2005.

[27] J. L. De La Vega-Beltran, C. Sánchez-Cárdenas, D. Krapf, E. O. Hernandez-González, E. Wertheimer, C. L. Treviño, P. E. Visconti, and A. Darszon. Mouse Sperm Membrane Potential Hyperpolarization Is Necessary and Sufficient to Prepare Sperm for the Acrosome Reaction. Journal of Biological Chemistry, 287(53):44384–44393, dec 2012.

[28] T. E. DeCoursey. Voltage-Gated Proton Channels: Molecular Biology, Physiology, and Pathophysiology of the HV Family. Physiological Reviews, 93(2):599–652, 2013.

[29] J. Espinal, M. Aldana, A. Guerrero, C. Wood, A. Darszon, and G. Martínez-Mekler. Discrete dynamics model for the speract-activated Ca 2+ signaling network relevant to sperm motility. PLoS ONE, 6(8):1–11, 2011.

[30] J. Espinal-Enríquez, D. A. Priego-Espinosa, A. Darszon, C. Beltrán, and G. Martínez-Mekler. Network model predicts that CatSper is the main Ca2+ channel in the regulation of sea urchin sperm motility. Scientific Reports, 7(1):4236, 2017.

[31] C. Espinosa-Soto, P. Padilla-Longoria, and E. R. Alvarez-buylla. A gene regulatory network model for cell-fate determination during Arabidopsis thalianal flower development that is robust and recovers experimental gene expression profiles. Plant Cell, 16(11):2923–2939, 2004.

[32] H. Florman and T. Ducibella. Fertilization in Mammals. In Knobil and Neill’s Physiology of Reproduction, pages 55–112. Elsevier, 2006.

[33] H. M. Florman, R. M. Tombes, N. L. First, and D. F. Babcock. An adhesion-associated agonist from the zona pellucida activates G protein-promoted elevations of internal Ca2+ and pH that mediate mammalian sperm acrosomal exocytosis. Developmental Biology, 1989.

[34] J. K. Foskett, C. White, K.-H. Cheung, and D.-O. D. Mak. Inositol trisphosphate receptor Ca2+ release channels. Physiological reviews, 87(2):593–658, apr 2007.

[35] K. Fukami, K. Nakao, T. Inoue, Y. Kataoka, M. Kurokawa, R. A. Fissore, K. Nakamura, M. Katsuki, K. Mikoshiba, N. Yoshida, and T. Takenawa. Requirement of phospholipase Cδ4 for the zona pellucida-induced acrosome reaction. Science, 2001.

[36] K. Fukami, M. Yoshida, T. Inoue, M. Kurokawa, R. A. Fissore, N. Yoshida, K. Mikoshiba, and T. Takenawa. Phospholipase Cδ4 is required for Ca2+ mobilization essential for acrosome reaction in sperm. Journal of Cell Biology, 161(1):79–88, 2003.

[37] B. Guyonnet, N. Egge, and G. A. Cornwall. Functional Amyloids in the Mouse Sperm Acrosome. Molecular and Cellular Biology, 34(14):2624–2634, 2014.

[38] C. Harper, L. Wootton, F. Michelangeli, L. Lefièvre, C. Barratt, and S. Publicover. Secretory pathway Ca2+-ATPase (SPCA1) Ca2+ pumps, not SERCAs, regulate complex [Ca2+]i signals in human spermatozoa. Journal of Cell Science, 118(8):1673–1685, apr 2005.

[39] C. V. Harper and S. J. Publicover. Reassessing the role of progesterone in fertilization—compartmentalized calcium signalling in human spermatozoa? Human Reproduction, 20(10):2675–2680, oct 2005.

[40] D. M. Hidalgo, A. Romarowski, M. G. Gervasi, F. Navarrete, M. Balbach, A. M. Salicioni, L. R. Levin, J. Buck, and P. E. Visconti. Capacitation increases glucose consumption in murine sperm. Molecular Reproduction and Development, 87(10):1037–1047, 2020.

[41] D. M. Hutt, J. M. Baltz, and J. K. Ngsee. Synaptotagmin VI and VIII and syntaxin 2 are essential for the mouse sperm acrosome reaction. Journal of Biological Chemistry, 280(21):20197–20203, 2005.

[42] D. M. Hutt, R. a. Cardullo, J. M. Baltz, and J. K. Ngsee. Synaptotagmin VIII is localized to the mouse sperm head and may function in acrosomal exocytosis. Biology of reproduction, 66(1):50–6, 2002.

[43] H. Iida, Y. Yoshinaga, S. Tanaka, K. Toshimori, and T. Mori. Identification of Rab3A GTPase as an acrosome-associated small GTP-binding protein in rat sperm. Developmental biology, 211:144–155, 1999.

[44] N. Inoue, Y. Satouh, M. Ikawa, M. Okabe, and R. Yanagimachi. Acrosome-reacted mouse spermatozoa recovered from the perivitelline space can fertilize other eggs. Proceedings of the National Academy of Sciences of the United States of America, 108(50):20008–11, 2011.

[45] M. Jin, E. Fujiwara, Y. Kakiuchi, M. Okabe, Y. Satouh, S. A. Baba, K. Chiba, and N. Hirohashi. Most fertilizing mouse spermatozoa begin their acrosome reaction before contact with the zona pellucida during in vitro fertilization. Proceedings of the National Academy of Sciences, 108(12):4892–4896, 2011.

[46] M. K. Jungnickel, K. A. Sutton, Y. Wang, and H. M. Florman. Phosphoinositide-dependent pathways in mouse sperm are regulated by egg ZP3 and drive the acrosome reaction. Developmental Biology, 304(1):116–126, 2007.

[47] S. A. Kauffman. Metabolic stability and epigenesis in randomly constructed genetic nets. Journal of Theoretical Biology, 22(3):437–467, 1969.

[48] Y. Kirichok, B. Navarro, and D. E. Clapham. Whole-cell patch-clamp measurements of spermatozoa reveal an alkaline-activated Ca2+ channel. Nature, 2006.

[49] S. Kleinboelting, A. Diaz, S. Moniot, J. Van Den Heuvel, M. Weyand, L. R. Levin, J. Buck, and C. Steegborn. Crystal structures of human soluble adenylyl cyclase reveal mechanisms of catalysis and of its activation through bicarbonate. Proceedings of the National Academy of Sciences of the United States of America, 111(10):3727–3732, mar 2014.

[50] C. Lawson, V. Dorval, S. Goupil, and P. Leclerc. Identification and localisation of SERCA 2 isoforms in mammalian sperm. MHR: Basic science of reproductive medicine, 13(5):307–316, may 2007.

[51] L. Lefièvre, K. Nash, S. Mansell, S. Costello, E. Punt, J. Correia, J. Morris, J. Kirkman-Brown, S. M. Wilson, C. L. R. Barratt, and S. Publicover. 2-APB-potentiated channels amplify CatSper-induced Ca(2+) signals in human sperm. The Biochemical journal, 448(2):189–200, 2012.

[52] P. Lishko, I. Botchkina, and Y. Kirichok. Progesterone activates the principal Ca2+ channel of human sperm Steroid signalling and ion channel regulation View project. Article in Nature, 2011.

[53] P. V. Lishko, I. L. Botchkina, A. Fedorenko, and Y. Kirichok. Acid Extrusion from Human Spermatozoa Is Mediated by Flagellar Voltage-Gated Proton Channel. Cell, 140(3):327–337, 2010.

[54] P. V. Lishko, Y. Kirichok, D. Ren, B. Navarro, J.-J. Chung, and D. E. Clapham. The Control of Male Fertility by Spermatozoan Ion Channels. Annual Review of Physiology, 2012.

[55] G. M. Luque, T. Dalotto-Moreno, D. Martín-Hidalgo, C. Ritagliati, L. C. Puga Molina, A. Romarowski, P. A. Balestrini, L. J. Schiavi-Ehrenhaus, N. Gilio, D. Krapf, P. E. Visconti, and M. G. Buffone. Only a subpopulation of mouse sperm displays a rapid increase in intracellular calcium during capacitation. Journal of Cellular Physiology, 2018.

[56] N. Mannowetz, M. R. Miller, P. V. Lishko, and D. E. Clapham. Regulation of the sperm calcium channel CatSper by endogenous steroids and plant triterpenoids. PNAS, 114(22):5743–5748, 2017.

[57] E. Mata-Martinez, A. Darszon, and C. L. Treviño. pH-dependent Ca+2 oscillations prevent untimely acrosome reaction in human sperm. Biochemical and Biophysical Research Communications, 497(1):146–152, feb 2018.

[58] L. S. Mayorga, C. N. Tomes, and S. A. Belmonte. Acrosomal exocytosis, a special type of regulated secretion. IUBMB Life, 59(4-5):286–292, 2007.

[59] S. Meizel, K. O. Turner, and R. Nuccitelli. Progesterone Triggers a Wave of Increased Free Calcium during the Human Sperm Acrosome Reaction. Developmental Biology, 182(1):67–75, 1997.

[60] L. Mendoza and E. R. Alvarez-Buylla. Dynamics of the genetic regulatory network for Arabidopsis thaliana flower morphogenesis. Journal of theoretical biology, 193(2):307–19, 1998.

[61] L. Mendoza and E. R. Alvarez-Buylla. Genetic regulation of root hair development in Arabidopsis thaliana: a network model. Journal of theoretical biology, 204(3):311–26, 2000.

[62] M. Michaut, C. N. Tomes, G. De Blas, R. Yunes, and L. S. Mayorga. Calcium-triggered acrosomal exocytosis in human spermatozoa requires the coordinated activation of Rab3A and N-ethylmaleimide-sensitive factor. Proceedings of the National Academy of Sciences, 97(18):9996–10001, 2000.

[63] M. Miller, S. Kenny, N. Mannowetz, S. Mansell, M. Wojcik, R. Zucker, K. Xu, and P. V. Lishko. Asymmetrically Positioned Flagellar Control Units Regulate Human Sperm Rotation. bioRxiv, 2018.

[64] M. R. Miller, N. Mannowetz, A. T. Iavarone, R. Safavi, E. O. Gracheva, J. F. Smith, R. Z. Hill, D. M. Bautista, Y. Kirichok, and P. V. Lishko. Unconventional endocannabinoid signaling governs sperm activation via the sex hormone progesterone. Science, 2016.

[65] L. C. Molina, S. Gunderson, J. Riley, P. Lybaert, A. Borrego-Alvarez, E. S. Jungheim, and C. M. Santi. Membrane Potential Determined by Flow Cytometry Predicts Fertilizing Ability of Human Sperm. Frontiers in Cell and Developmental Biology, 7(January):1–12, 2020.

[66] T. Murase and E. R. S. Roldan. Progesterone and the zona pellucida activate different transducing pathways in the sequence of events leading to diacylglycerol generation during mouse sperm acrosomal exocytosis. Biochemical Journal, 320(3):1017–1023, dec 1996.

[67] R. A. Nahed, G. Martinez, J. Escoffier, S. Yassine, T. Karaouzène, J. P. Hograindleur, J. Turk, G. Kokotos, P. F. Ray, S. Bottari, G. Lambeau, S. Hennebicq, and C. Arnoult. Progesterone-induced acrosome exocytosis requires sequential involvement of calcium-independent phospholipase A2β (iPLA2β) and group X secreted phospholipase A2(sPLA2). Journal of Biological Chemistry, 291(6):3076–3089, 2016.

[68] N. Nakamura, S. Tanaka, Y. Teko, K. Mitsui, and H. Kanazawa. Four Na+/H+ exchanger isoforms are distributed to Golgi and post-Golgi compartments and are involved in organelle pH regulation. Journal of Biological Chemistry, 280(2):1561–1572, 2005.

[69] T. Nakanishi, M. Ikawa, S. Yamada, K. Toshimori, and M. Okabe. Alkalinization of Acrosome Measured by GFP as a pH Indicator and Its Relation to Sperm Capacitation. Developmental Biology, 237(1):222–231, 2001.

[70] T. Nishigaki, O. José, A. L. González-Cota, F. Romero, C. L. Treviño, and A. Darszon. Intracellular pH in sperm physiology. Biochemical and Biophysical Research Communications, 450(3):1149–1158, aug 2014.

[71] K. Oberheide, D. Puchkov, and T. J. Jentsch. Loss of the Na+/H+ exchanger NHE8 causes male infertility in mice by disrupting acrosome formation. Journal of Biological Chemistry, 292(26):10845–10854, 2017.

[72] M. Okabe. The Acrosome Reaction: A Historical Perspective. In Sperm Acrosome Biogenesis and Function During Fertilization, pages 1–13. Springer, Cham, 2016.

[73] N. Okamura, Y. Tajima, A. Soejima, H. Masuda, and Y. Sugita. Sodium bicarbonate in seminal plasma stimulates the motility of mammalian spermatozoa through direct activation of adenylate cyclase. Journal of Biological Chemistry, 260(17):9699–9705, 1985.

[74] G. Orta, J. L. de la Vega-Beltran, D. Martín-Hidalgo, C. M. Santi, P. E. Visconti, and A. Darszon. CatSper channels are regulated by protein kinase A. Journal of Biological Chemistry, 293(43):16830–16841, oct 2018.

[75] J. Ramalho-Santos, R. D. Moreno, P. Sutovsky, a. W. Chan, L. Hewitson, G. M. Wessel, C. R. Simerly, and G. Schatten. SNAREs in mammalian sperm: possible implications for fertilization. Developmental biology, 223(1):54–69, 2000.

[76] J. Ramalho-Santos and G. Schatten. Presence of N-ethyl maleimide sensitive factor (NSF) on the acrosome of mammalian sperm. Systems Biology in Reproductive Medicine, 50(3):163–168, 2004.

[77] P. Redecker, M. R. Kreutz, J. Bockmann, E. D. Gundelfinger, and T. M. Boeckers. Brain synaptic junctional proteins at the acrosome of rat testicular germ cells. The journal of histochemistry and cytochemistry: official journal of the Histochemistry Society, 51(6):809–19, 2003.

[78] M. F. Romero, A. P. Chen, M. D. Parker, and W. F. Boron. The SLC4 family of bicarbonate (HCO3-) transporters, 2013.

[79] M. Rossato, F. Di Virgilio, C. Galeazzi, and C. Foresta. Intracellular calcium store depletion and acrosome reaction in human spermatozoa: Role of calcium and plasma membrane potential. Molecular Human Reproduction, 7(2):119–128, 2001.

[80] C. Sánchez-Cárdenas, M. R. Servín-Vences, O. José, C. L. Treviño, A. Hernández-Cruz, and A. Darszon. Acrosome Reaction and Ca2+ Imaging in Single Human Spermatozoa: New Regulatory Roles of [Ca2+]i1. Biology of Reproduction, 91(3):1–13, 2014.

[81] C. Schmeitz, E. A. Hernandez-Vargas, R. Fliegert, A. H. Guse, and M. Meyer-Hermann. A mathematical model of T lymphocyte calcium dynamics derived from single transmembrane protein properties. Frontiers in Immunology, 4(SEP), 2013.

[82] K. Schuh, E. J. Cartwright, E. Jankevics, K. Bundschu, J. Liebermann, J. C. Williams, A. L. Armesilla, M. Emerson, D. Oceandy, K. P. Knobeloch, and L. Neyses. Plasma membrane Ca2+ ATPase 4 is required for sperm motility and male fertility. Journal of Biological Chemistry, 279(27):28220–28226, 2004.

[83] J. R. Schulz, J. D. Sasaki, and V. D. Vacquier. Increased association of synaptosome-associated protein of 25 kDa with syntaxin and vesicle-associated membrane protein following acrosomal exocytosis of sea urchin sperm. Journal of Biological Chemistry, 273(38):24355–24359, 1998.

[84] C. Soriano-Úbeda, J. Romero-Aguirregomezcorta, C. Matás, P. E. Visconti, and F. A. García-Vázquez. Manipulation of bicarbonate concentration in sperm capacitation media improves in vitro fertilisation output in porcine species. Journal of Animal Science and Biotechnology, 10(19), 2019.

[85] C. M. Sosa, M. A. Pavarotti, M. N. Zanetti, F. C. M. Zoppino, G. A. De Blas, and L. S. Mayorga. Kinetics of human sperm acrosomal exocytosis. Molecular Human Reproduction, 21(3):244–254, 2014.

[86] C. M. Sosa, M. N. Zanetti, C. A. Pocognoni, and L. S. Mayorga. Acrosomal Swelling Is Triggered by cAMP Downstream of the Opening of Store-Operated Calcium Channels During Acrosomal Exocytosis in Human Sperm. Biology of reproduction, 94(3):57, 2016.

[87] C. Steegborn. Structure, mechanism, and regulation of soluble adenylyl cyclases - similarities and differences to transmembrane adenylyl cyclases, dec 2014.

[88] C. Stival, L. D. C. Puga Molina, B. Paudel, M. G. Buffone, P. E. Visconti, and D. Krapf. Sperm Capacitation and Acrosome Reaction in Mammalian Sperm. Advances in anatomy, embryology, and cell biology, 220:93–106, 2016.

[89] T. Strünker, N. Goodwin, C. Brenker, N. D. Kashikar, I. Weyand, R. Seifert, and U. B. Kaupp. The CatSper channel mediates progesterone-induced Ca2+ influx in human sperm. Nature, 471(7338):382–387, 2011.

[90] G.-H. Sun-Wada, Y. Imai-Senga, A. Yamamoto, Y. Murata, T. Hirata, Y. Wada, and M. Futai. A Proton Pump ATPase with Testis-specific E1-Subunit Isoform Required for Acrosome Acidification. THE JOURNAL OF BIOLOGICAL CHEMISTRY, 277(20):18098–18105, 2002.

[91] M. E. Teves, H. A. Guidobaldi, D. R. Uñates, R. Sanchez, W. Miska, S. J. Publicover, A. A. Morales Garcia, and L. C. Giojalas. Molecular Mechanism for Human Sperm Chemotaxis Mediated by Progesterone. PLoS ONE, 4(12):e8211, dec 2009.

[92] D. Thieffry. Dynamical roles of biological regulatory circuits. Briefings in Bioinformatics, 2007.

[93] C. N. Tomes. The proteins of exocytosis: lessons from the sperm model. The Biochemical journal, 465(3):359–70, 2015.

[94] C. N. Tomes, G. A. De Blas, M. A. Michaut, E. V. Farré, O. Cherhitin, P. E. Visconti, and L. S. Mayorga. α-SNAP and NSF are required in a priming step during the human sperm acrosome reaction. Molecular Human Reproduction, 11(1):43–51, 2005.

[95] C. N. Tomes, M. Michaut, G. De Blas, P. Visconti, U. Matti, and L. S. Mayorga. SNARE complex assembly is required for human sperm acrosome reaction. Developmental biology, 243(2):326–338, 2002.

[96] R. Trejo and A. Mújica. Changes in calmodulin compartmentalization throughout capacitation and acrosome reaction in guinea pig spermatozoa. Molecular Reproduction and Development, 26(4):366–376, 1990.

[97] P. E. Visconti, G. D. Moore, J. L. Bailey, P. Leclerc, S. A. Connors, D. Pan, P. Olds-clarke, and G. S. Kopf. Capacitation of mouse spermatozoa. Development, 1150:1139–1150, 1995.

[98] P. E. Visconti, V. A. Westbrook, O. Chertihin, I. Demarco, S. Sleight, and A. B. Diekman. Novel signaling pathways involved in sperm acquisition of fertilizing capacity. Journal of Reproductive Immunology, 53(1-2):133–150, 2002.

[99] D. Wang, J. Hu, I. A. Bobulescu, T. a. Quill, P. McLeroy, O. W. Moe, and D. L. Garbers. A sperm-specific Na+/H+ exchanger (sNHE) is critical for expression and in vivo bicarbonate regulation of the soluble adenylyl cyclase (sAC). Proceedings of the National Academy of Sciences of the United States of America, 104(22):9325–9330, 2007.

[100] C. R. Ward, D. Faundes, and J. A. Foster. The monomeric GTP binding protein, rab3a, is associated with the acrosome in mouse sperm. Molecular Reproduction and Development, 53(4):413–421, 1999.

[101] P. M. Wassarman and E. S. Litscher. The multifunctional zona pellucida and mammalian fertilization. Journal of Reproductive Immunology, 83(1-2):45–49, 2009.

[102] A. I. Yudin, W. Gottlieb, and S. Meizel. Ultrastructural studies of the early events of the human sperm acrosome reaction as initiated by human follicular fluid. Gamete research, 20(1):11–24, 1988.

[103] R. Yunes, M. Michaut, C. Tomes, and L. S. Mayorga. Rab3A Triggers the Acrosome Reaction in Permeabilized Human Spermatozoa. BIOLOGY OF REPRODUCTION, 62:1084–1089, 2000.

[104] D. Zhang and M. Gopalakrishnan. Sperm ion channels: Molecular targets for the next generation of contraceptive medicines? Journal of Andrology, 26(6):643–653, 2005.

[105] X. Zhang, X. Zeng, and C. J. Lingle. Slo3 K+ Channels: Voltage and pH Dependence of Macroscopic Currents. The Journal of General Physiology, 128(3):317–336, 2006.

[106] L. Zhao, X. Shi, L. Li, and D. J. Miller. Dynamin 2 associates with complexins and is found in the acrosomal region of mammalian sperm. Molecular Reproduction and Development, 74(6):750–757, 2007.

