## Appendix A for "Discrete dynamic model of the mammalian sperm acrosome reaction: the influence of acrosomal pH and biochemical heterogeneity"

### Appendix A: Construction of $IP_3R_a$ regulatory function as an example

Acrosomal $IP_3$ receptors ($IP_3R_a$) are $[Ca^{2+}]_i$ and $IP_3$ dependent $Ca^{2+}$ channels located on the outer acrosomal membrane. They posses one $IP_3$ binding site and two $Ca^{2+}$ binding sites of high and low affinities that open or block the channel respectively. Increasing $[Ca^{2+}]_i$ in presence of $IP_3$ promotes opening of the channel. A further increase in $[Ca^{2+}]_i$ blocks the channel by means of the low affinity $Ca^{2+}$ binding site.

Table S1 presents the regulatory function of the $IP_3R_a$. For its construction, we first identified the nodes $IP_3$ and $[Ca^{2+}]_i$ as its regulators. $IP_3$ can take only two values ( Basal=0 and Increased=1 ) while $[Ca^{2+}]_i$ can take three values (Basal=0, Activator=1, Inhibitor=2). $[Ca^{2+}]_i$=1 represents an increase in $[Ca^{2+}]_i$ sufficient to open the $IP_3R_a$ in presence of an $IP_3$ increment while $[Ca^{2+}]_i$=2 promotes its blockade. For practical purposes, the $IP_3R_a$ can take one of two possible values (Open=1 and Closed=0), depending entirely on the value of its regulators. When $IP_3$=0, the channel can not open independently of the value of $[Ca^{2+}]_i$. When $[Ca^{2+}]_i$=0 or $[Ca^{2+}]_i$=2, the channel is closed because $[Ca^{2+}]_i$ is either to low or to high, independently of the value of $IP_3$. This means that $IP_3R_a$=1 only when $IP_3$=1 and $[Ca^{2+}]_i$=1. The value of $IP_3R_a$ and in general of every node should be specified for every possible value of its regulators.

| **Regulators** | | **Target** |
|---|---|---|
| **$IP_3$** | **$[Ca^{2+}]_i$** | **$IP_3R_a$** |
| 0 | 0 | 0 |
| 0 | 1 | 0 |
| 0 | 2 | 0 |
| 1 | 0 | 0 |
| 1 | 1 | 1 |
| 1 | 2 | 0 |

Table S1: $IP_3R_a$ regulatory function. Rows represent the value assigned to $IP_3R_a$ under each possible value of its regulators $IP_3$ and $[Ca^{2+}]_i$
